# The mutational load in natural populations is significantly affected by high primary rates of retroposition

**DOI:** 10.1101/2020.08.06.239277

**Authors:** Wenyu Zhang, Chen Xie, Kristian Ullrich, Yong E. Zhang, Diethard Tautz

## Abstract

Gene retroposition is known to contribute to patterns of gene evolution and adaptations. However, possible negative effects of gene retroposition remain largely unexplored, since most previous studies have focussed on between-species comparisons where negatively selected copies are mostly not observed, as they are quickly lost from the populations. Here, we show for natural house mouse populations that the primary rate of retroposition is orders of magnitude higher than previously thought. Comparisons with SNP distribution patterns in the same populations show that most retroposition events are deleterious. Transcriptomic profiling analysis shows that new retroposed copies become easily subject to transcription and have an influence on the expression level of their parental genes, especially when transcribed in the antisense direction. Our results imply that the impact of retroposition on the mutational load in natural populations has been highly underestimated, which has also implications for strategies of disease allele detection in humans.

**Significance statement:** The phenomenon or retroposition (re-integration of reverse transcribed RNA into the genome), has been well studied in comparisons between genomes and has been identified as a source of evolutionary innovation. However, the negative effects of retroposition have been overlooked so far. Our study makes use of a unique population genomic dataset from natural mouse populations. It shows that the retroposition rate is magnitudes higher than previously suspected. We show that most of the newly transposed retrocopies have a deleterious impact through modifying the expression of their parental genes. In humans, this effect is expected to cause disease alleles and we propose that genetic screening needs to take into account the search for newly transposed retrocopies.

## Introduction

Gene retroposition (or RNA-based gene duplication) is a particular type of gene duplication in which a gene’s transcript is used as a template to generate new gene copies (retrocopies), and this has a variety of evolutionary implications (1–3). The intronless retrocopies have initially been viewed as evolutionary dead-ends with little biological effects (4, 5), mainly due to the assumed lack of regulatory elements and promotors. However, this hypothesis has become of less relevance after it has become clear that a large portion (>80%) of the mammalian genome is transcribed (6, 7) and that there is a fast evolutionary turnover of these transcribed regions, implying that essentially every part of the genome is accessible to transcription (8). In addition, retrocopies can recruit their own regulatory elements through a number of mechanisms (2, 3). Hence, retrocopies can act as functional retrogenes that encode full-length proteins, and it has, therefore, been proposed that they contribute to the evolution of new biological functions through neo-functionalization or sub-functionalization (2, 3, 9–11). However, the possibility that retroposition events could also be deleterious has been much less considered so far. Deleterious effects could be due to insertions into functional sites, and this has indeed been detected in a retrogene population analysis in humans (12). However, even if they land in nonfunctional intergenic regions, they could still be transcribed and their transcripts could interfere with the function of the parental genes (13–15). In SNP based association studies, this would become apparent as a trans-effect on the parental gene, but the true reason for the trans-effect would remain unnoticed when the retrocopy is not included in the respective genomic reference sequence. Hence, if transposition rates are high and if the transposed copies are frequently transcribed, they could have a substantial impact on the mutational landscape of genomes.

Retroposition mechanisms were initially studied in between-species comparisons with single genomes per species (*e.g*. (16)), but these will miss all cases of retropositions with deleterious effects. Accumulating population genomics data are now providing the opportunity to detect novel retroposed gene copy number variants (retroCNVs) that are still polymorphic in populations (3), but a broad comparative dataset from related evolutionary lineages is required to obtain a deeper insight. A population analysis representing natural samples is available in humans, based on the 1,000 Genomes Project Consortium data (12, 17–20). However, the power of the discovery of retroCNVs in these studies has been limited due to the heterozygous and relatively low coverage sequencing datasets. Moreover, in humans it is not possible to compare the data with very closely related other lineages, since these are extinct (*e.g*., Neandertals or Denisovans). Hence, a comprehensive analysis on the evolutionary dynamics of retroCNVs at comparable individual genome level, especially based on a set of well-defined natural populations from different lineages where evolutionary processes and transposition rates can be studied, is still missing.

The house mouse (*Mus musculus*) is a particularly suitable model system for comparative genomic analyses in natural populations, owing to its well-studied evolutionary history (21, 22). Currently, three major lineages of *Mus musculus* are distinguished, classified as subspecies which diverged roughly 0.5 million year ago: the western house mouse *Mus musculus domesticus*, the eastern house mouse *Mus musculus musculus* and the southeast-Asian house mouse *Mus musculus castaneus*. Previously, we have generated a unique genomic resource using wild mice collected from multiple geographic regions (each representing one natural population) covering these three major house mouse subspecies, with a carefully designed sampling procedure to maximize the possibility of capturing the genetic diversity from each population (23). This was complemented by a well-controlled experimental set-up to generate largely homogeneous genomic/transcriptomic sequencing datasets at relatively high coverage for the same individuals (24), which makes it possible to trace directly the effects of new retroposed copies on the expression of their parental genes.

Here we show that the turnover (gain and loss) rates of retroCNVs are many-fold higher than previously expected and the frequency spectra of retroCNV alleles in populations in comparison to SNP allele frequency spectra implies mostly deleterious effects. Transcriptome data show that the new retroCNVs are usually transcribed and have indeed an effect on the parental gene transcripts. A new strand-specific RNA-Seq dataset for one of the populations shows that antisense transcribed retroCNVs are highly underrepresented compared to sense transcripts, implying strong selection against them. We conclude that deleterious effects of newly transposed retrocopies of genes have been largely underestimated so far. We also discuss the implications for human disease allele detection.

## Results

Full genome resequencing data of 96 individuals derived from nine natural populations corresponding to the three major subspecies (*M. m. domesticus, M. m. musculus and M. m. castaneus*) of the house mouse (*Mus musculus*), as well as nine individuals from two outgroup species (*M. spicilegus* and *M. spretus*), were used to assess gene retroposition events (Fig. 1 and *SI Appendix*, Tables S1 and Dataset S1A). By adapting an exon-exon junction and exon-intron-exon junction mapping based approach for short read genomic sequencing data (18, 19, 25), we refined a computational pipeline to identify retroposition events (*SI Appendix*, Text S1) including a power analysis for optimizing mapping conditions. A retroposition event is identified, on condition that both the intron loss and the presence of a parental gene can be observed in the same individual sequencing dataset (25).

**Fig. 1:**
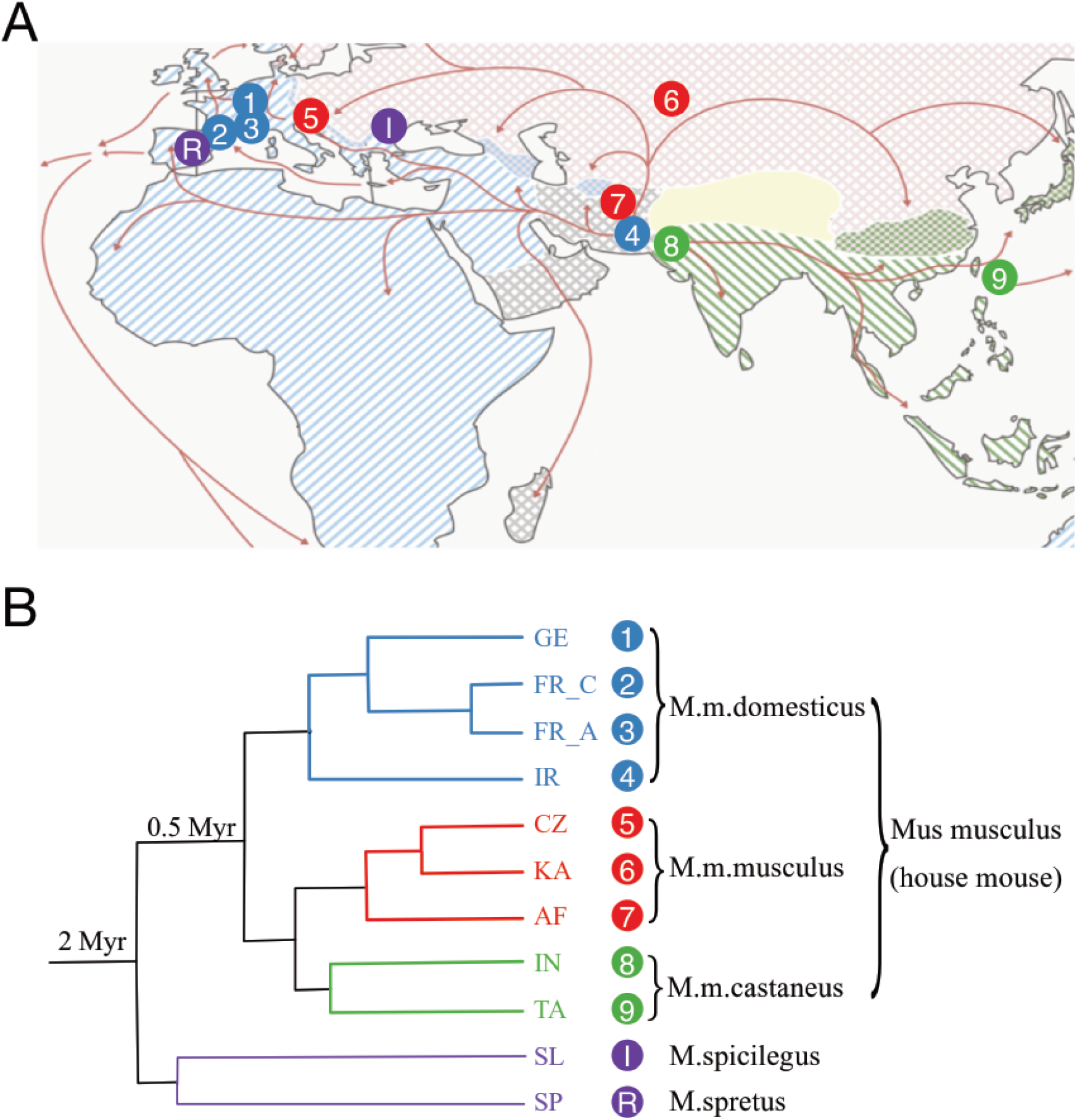
Geographic locations and phylogenetic relationships of house mouse subspecies and two out-group species samples. (A) Geographic location information on the sampled mouse individuals. This map is modified from (24). Territory areas for each house mouse subspecies: *M. m. domesticus* (blue); *M. m. musculus* (red); *M. m. castaneus* (green). Red arrows indicate possible migration routes, mostly during the spread of agriculture and trading. Geographic locations: 1, Cologne-Bonn/Germany (GE); 2, Massif Cenral/France (FR_C); 3, Auvergne-Rhône-Alpes/France (FR_A); 4, Ahvaz/Iran (IR); 5, Studenec/Czech Republic (CZ); 6, Almaty/Kazakhstan (KA); 7, Mazar/Afghanistan (AF); 8, Himachal Pradesh/India (IN); 9, Taiwan (TA); I, Sása/Slovakia (SL); R, Madrid/Spain (SP). (B) Phylogenetic relationships and split time estimates (branches not shown to scale) among the house mouse populations and two out-group species in the study.

Due to the need to detect at least one exon-exon junction, only protein coding genes with ≥ 2 exons (~ 92.4% of all coding genes annotated in Ensembl v87) were assayed as potential source of gene retroposition. To compensate for the variance in the sequencing (read length, coverage, and etc.) and individual intrinsic features (*i.e*., sequence divergence from the mm10 reference genome), we optimized the parameters (*i.e*., alignment identity, spanning read length, and number of supporting reads) of the retroposition event discovery pipeline for each individual genome (*SI Appendix*, Text S1). The resultant computational pipeline gave a low false positive discovery rate < 3% (*SI Appendix*, Fig. S2) and a high recall rate of > 95% (*SI Appendix*, Fig. S5) for all the individual genomes tested. This optimization ensures that the calling probability for retroposition events is in the same order as that for SNP calling based on GATK (26), *i.e*., retroCNV and SNP frequency data become comparable.

A subset of the retroCNV alleles that were identified as newly arisen in one of our populations is also present in the mm10 reference genome. For these ones, we directly called their presence based on the alignment data of individual sequencing datasets to the reference genome. For those retroCNV alleles that are absent in the mm 10 reference genome, we inferred their insertion sites in the genome by using discordant aligned paired-end reads when these could be uniquely mapped (see Methods). Additionally, a detailed discussion on the possible technical issues of retroCNV discovery can be found in *SI Appendix* Text S2.

### High numbers of retroCNVs in natural populations

Applying the above pipeline, we screened for retroCNV parental genes (*i.e*., the parental genes from which retrocopies are derived) and retroCNVs (*i.e*., alleles of the inserted retrocopies, or insertion sites in the genome in case that the retrocopies are not present in the reference genome) in the mouse individual genome sequencing datasets. To study turnover rates (*i.e*., gains and losses), We focused on the recently originated gene retroposition events in the house mouse lineage, *i.e*., retroCNV parental genes and retroCNVs occurring in the *Mus musculus* subspecies but absent in the outgroup species.

In total, we identified 21,160 house mouse (*Mus musculus*) specific retroposition events across all 96 individuals surveyed (*SI Appendix*, Fig. S6), whereby this number includes also those detected in more than one individual, as well as 8,483 for which no insertion site could be mapped (note that we omitted these from the more detailed analysis below). They are derived from 1,663 unique retroCNV parental genes (Dataset S2). Only 80 (4.8%) of these retroCNV parental genes have annotated recently originated retrocopies in the mm10 reference genome (≥ 95% alignment identity to their parental gene) based on RetrogeneDB v2 (27), while the other 1,583 retroCNV parental genes represent newly detected gene retroposition events in house mouse wild individuals. Approximately 3.9% of them show more than one retroCNV allele for the same retroCNV parental gene in the same individual genome (*SI Appendix*, Fig. S8).

Random resampling analysis of subsamples of individuals showed that the number of detectable retroCNV parental genes has not reached saturation with the given number of sampled individuals in our dataset (Fig. 2A), implying that many more retroCNV transposition events should be found when more individuals would be analyzed. Importantly, as suggested by (28), we also found that CNV detection pipelines that do not specifically consider retroCNVs, underestimate their prevalence. In a direct comparison with data from genic CNV detection (29), only <1%, on average, of the retroCNV parental genes detected in our analysis overlaps with genic CNVs according to this pipeline (*SI Appendix*, Fig. S9).

**Fig. 2:**
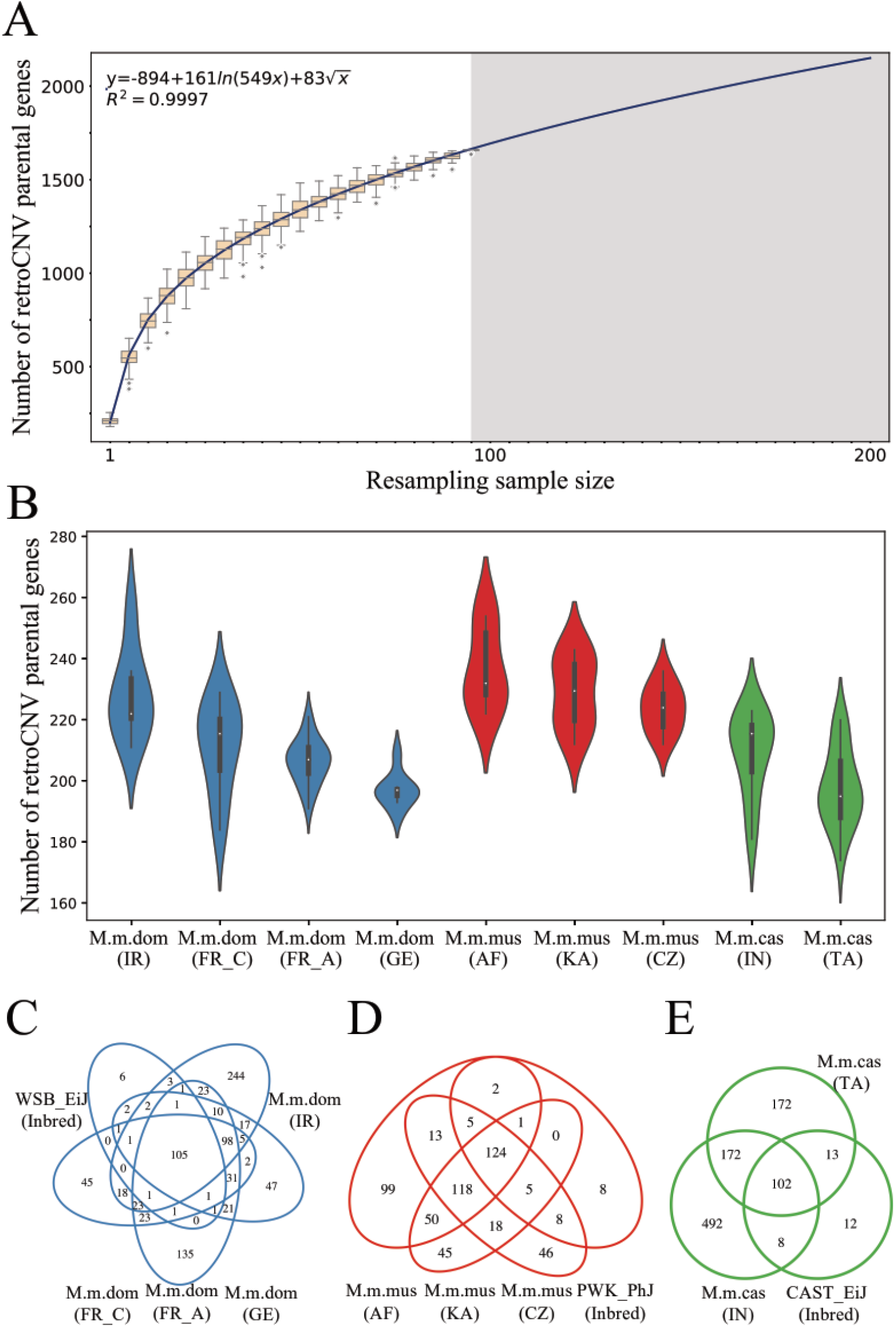
Distribution of the number of detected retroCNV parental genes across house mouse populations. Only *Mus musculus* specific retroCNV parental genes are included in this analysis. (A) Number of detected retroCNV parental genes with various random resampling sample sizes. The resampling subsample sizes were selected from 1 to 95, with step size of 5. Data points represent the average number of detected retroCNV parental genes of 100 replicates for each subsample, and whiskers the standard variance of the mean deviation. The gray area shows the prediction after doubling the number of current sampling house mouse individuals. (B) Distribution of the number of detected retroCNV parental genes within each house mouse natural population (see *SI Appendix*, Fig. S10 for a corresponding depiction of retroCNVs). (C) – (E) Depiction of the overlap of detected retroCNV parental genes between house mouse natural populations and inbred mouse lines derived from each of the three house mouse subspecies, respectively. Inbred mouse strains for three subspecies: WSB_EiJ (*M. m. domesticus*); PWK_PhJ (*M. m. musculus*); CAST_EiJ (*M. m. castaneus*). Abbreviations for geographic regions: *IR*, Iran; *FR_C*, France (Massif Central); *FR_A*, France (Auvergne-Rhône-Alpes); *GE*, Germany; *AF*, Afghanistan; *KA*, Kazakhstan; *CZ*, Czech Republic; IN, India; *TA*, Taiwan (see also Fig. 1A for geographic representation).

On average, in each tested individual, there are 212 retroCNV parental genes, but the populations differ somewhat in these numbers (Fig. 2B). Slightly higher numbers were found in the ancestral populations (*i.e*., Iran population for *M. m. domesticus*, Afghanistan population for *M. m. musculus*, and India population for*M. m. castaneus*), presumably since they have higher effective population sizes where more neutral or nearly neutral retroCNVs could segregate. The majority of retroCNV parental genes (91% - 95%) in the wild-derived laboratory inbred strains representing the three subspecies (*M. m. domesticus*: WSB_EiJ; *M. m. musculus*: PWK_PhJ; *M. m. castaneus*: CAST_Eij) can also be discovered in house mouse wild individuals (Fig. 2C-E). Conversely, the majority of retroCNV parental genes (73% - 87%) in wild-derived house mouse individuals are not present in the inbred mouse strains, since these represent essentially only single haplotypes from the wild diversity.

Among the above detected retroposition events for wild house mouse individuals, between 38% - 78% of their insertion sites in the genome could be identified (*SI Appendix*, Fig. S10), depending on the nature of the sequencing read data features of each individual, *e.g*., sequencing coverage, read length and insert size. The detection rate of insertion sites at the individual genome level presented here is much higher than the one that was reported from pooled human population genomes when the same criteria to define reliable insertion sites where applied (30% in (12)). Following the “gold standard” for calling novel retrocopies (*i.e*., with detectable genomic insertions, (20)), unless stated separately, all the following analysis were conducted on the basis of retroCNVs (corresponding to 12,677 retroposition events with detected insertion site), rather than retroCNV parental genes. Correspondingly, we included 2,025 unique house mouse specific retroCNVs (after collapsing of the same retroCNV alleles detected in multiple house mouse individuals, see Materials and Methods and *SI Appendix*, Fig. S6) for the further analysis (Dataset S3). Note that reliable SNP calling depends also on the need for unique mapping of reads, *i.e*., the reduced set is directly comparable to high quality SNP data.

### Rapid loss of retroCNVs

With SNP calling data from the same set of house mouse wild-derived individuals (see Methods), we were able to explore the retroCNV variation at different levels, in direct comparison to the SNP variation. For both retroCNV and SNP alleles, the frequency was computed by counting individuals with positive evidence of each allele, without distinguishing the homozygous and heterozygous genotype status. If one assumes that the SNPs are mostly neutral, they can be used as expectation for the demographic drift effects in the dataset. Of the 76,882,435 house mouse specific SNPs, 16.3% are found in all three house mouse subspecies (Fig. 3A), about 11% segregate in all nine populations (Fig. 3B) and 6.6% are found in all 96 tested house mouse individuals (Fig. 3C). Among the entire 2,025 different house mouse specific retroCNV alleles with mapped insertion site (Dataset S3), only 71 (3.5%) are found in all three house mouse subspecies (Fig. 3A) and only about 1% segregate in all nine populations (Fig. 3B), while none are found in all tested house mouse individuals (Fig. 3C). An additional analysis by using a subset of 1,551 retroCNVs (Dataset S3) that show both the positive evidence of retroCNV presence (*i.e*., detectable retroCNV allele) and the positive evidence of retroCNV absence (*i.e*., reliable alignments to span the retroCNV allele) in all 96 tested house mouse individuals (See Materials and Methods), confirmed the same observation that retroCNVs are more skewed toward singletons than are SNPs (*SI Appendix*, Fig. S11). This suggests that retroCNVs are removed not only through drift, but also through negative selection in the different lineages. This selective purging has the effect that the prevalence of retroposition rates will be underestimated when compared at the species or subspecies level only. In the following, we provide therefore an estimate for the most recent population splits in our dataset.

**Fig. 3:**
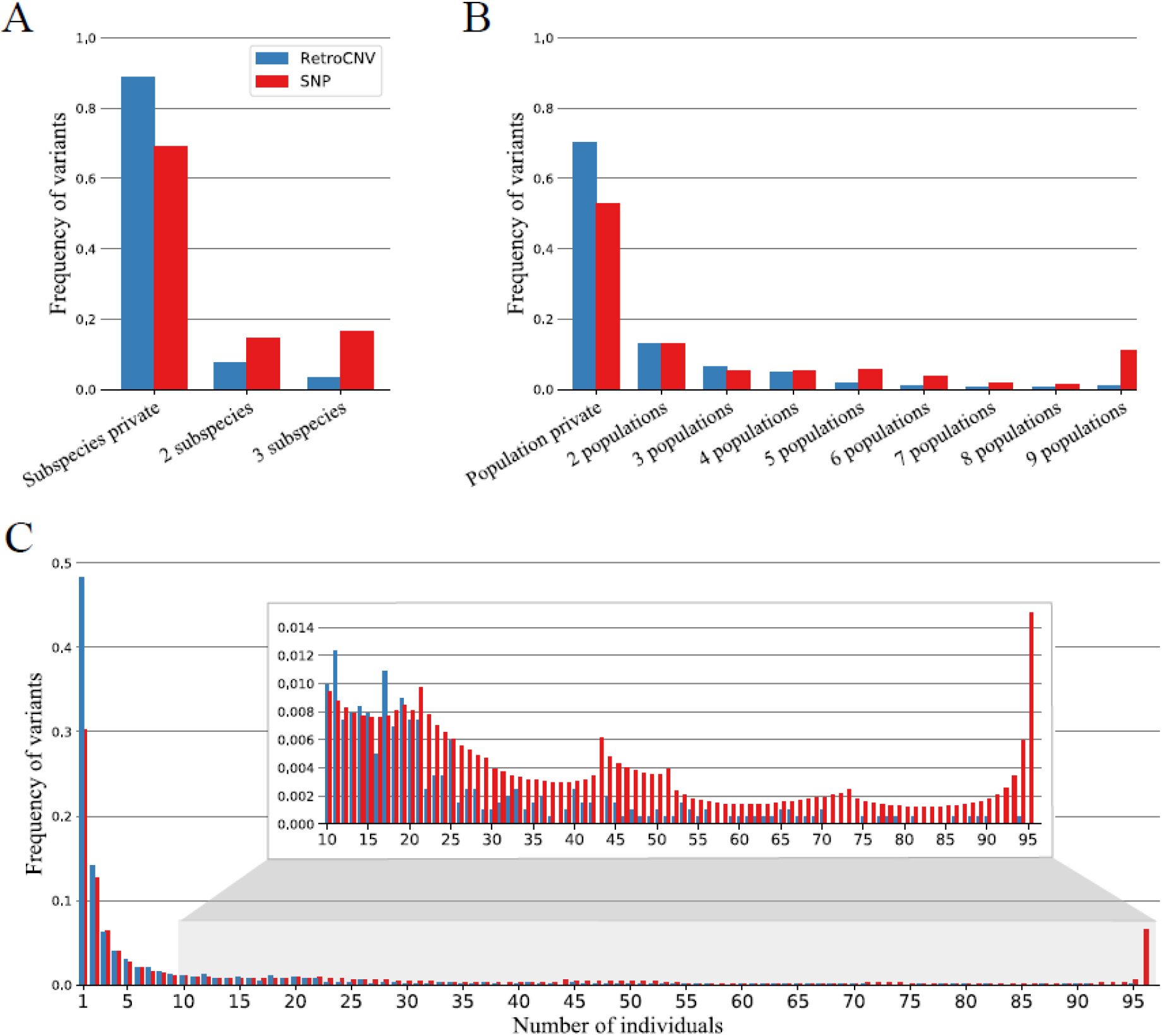
Distribution of the frequency of detected retroCNVs with mapped insertion sites and SNPs across different house mouse subspecies (A), populations (B) and individuals (C). The enlarged inset box in (C) is focused on the frequencies of retroCNVs/SNPs present in larger number of individuals.

The Western European *M. m. domesticus* populations are derived from Iranian (IR) populations and invaded Western Europe about 3,000 years ago where they quickly radiated. The split from the Iranian population would have occurred no more than 10,000 years ago (30, 31). This provides a time line to estimate retroCNV emergence rates by comparing the population and lineage-specific retroCNVs, under the assumption that they represent mostly new retroposition events in their lineage. We used the populations FR_C, GE and IR for this, since they are represented by the same number of individuals and were sequenced in a similar way. We found 60 and 57 private retroCNVs in FR_C and GE, respectively (Dataset S3). Assuming these populations have split soon after their arrival, this would suggest in the order of 200 new retroCNV events in 10,000 years. In the IR population, we find 284 private retroCNVs (Dataset S3), *i.e*., assuming a separation of 10,000 years, this would be of the same order.

A systematic comparison between primate species had suggested an birth rate of 21 to 160 retrocopies per million years (16), *i.e*., we estimate an about two orders of magnitude higher primary rate in our data, due to looking at a recent split, as well as population samples rather than single individuals. Indeed, when one increases the population sample, one can find even more population-specific retroCNVs, as becomes evident in the comparison between FR_C (N = 8) with FR_A (N = 20), where we found 60 versus 136 population-specific retroCNVs (Dataset S3). Hence, the number of primary retroposition events could even be higher, which explains also why we do not reach saturation of retroCNV parental genes even in our full sample set (Fig. 2A).

Negative selection effects can also be detected in the site frequency spectra analysis of the retroCNVs (Fig. 4) in comparison to the corresponding frequency spectra of SNP allele categories for the same population samples. Based on the functions of these SNPs, we categorized them into four distinct groups (32): 1) High effect SNPs that change the coding gene structure (stop codons or splice sites); 2) Moderate effect SNPs that change amino acid sites; 3) Low effect SNPs with synonymous changes; 4) Modifier effect SNPs that locate in non-coding regions. We found significantly more retroCNVs in the private category, *i.e*., occurring only in a single animal for each of the categories (Fisher’s exact test, retroCNV vs. high effect SNPs: p-value = 1.7 x 10^-18^; retroCNV vs. moderate effect SNPs: p-value = 2.6 x 10^-18^; retroCNV vs. low effect SNPs: p-value = 3.5 x 10^-67^; retroCNV vs. modifier effect SNPs: p-value = 1.3 x 10^-64^).

**Fig. 4.**
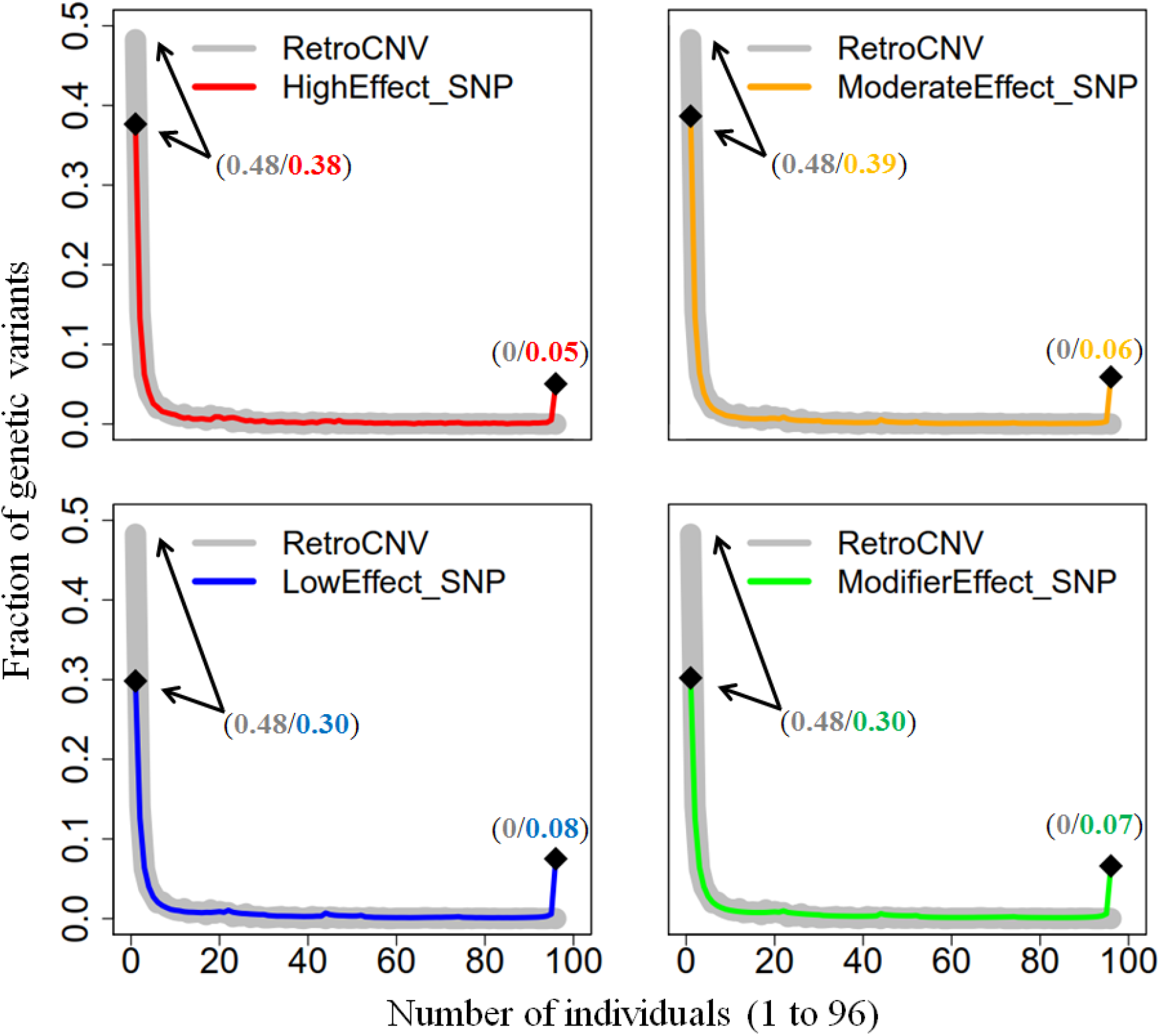
Comparison of the frequency spectrum of retroCNVs with the site frequency spectra of SNPs. High effect SNPs: the ones causing the gain/loss of start/stop codon or change the splicing acceptor/donor sites; Moderate effect SNPs: the ones resulting in a different amino acid sequence; Low effect SNPs: the ones occurring within the general region of the splice site, changing the final codon of an incompletely annotated transcript, changing the bases of start/stop codon (while start/terminator remains), or where there is no resulting change to the encoded amino acid; Modifier effect SNPs: the ones occurring around the coding regions of the genes (UTR, intron, up/downstream), noncoding gene regions, or intergenic regions. The numbers within the parentheses indicate the fractions of retroCNVs (in grey) and SNPs (colors corresponding to SNP categories) that are individual private or reach fixation in all 96 tested house mouse individuals, respectively.

To test for similarity of the distributions, we used the two sided Kolmogorov-Smirnov tests and found more similar distributions between retroCNVs and the more constrained SNP categories (Kolmogorov’s D statistic for retroCNV vs. high effect SNPs: D=0.14, retroCNV vs. moderate effect SNPs: D=0.13, retroCNV vs. low effect SNPs: D=0.21, retroCNV vs. modifier effect SNPs: D=0.21). Hence, from these data we conclude also that most new retroCNVs are under negative selection, *i.e*., would not only be lost by drift, but also largely lost by selective purging in natural populations.

### RetroCNV expression

In a previous study on the evolutionary origin of promotors of retrocopies (33), it was found that most retrocopies show at least a low level transcription whereby only about 3% of them inherited the promotor from the parental gene, while the remainder recruited it from a gene in the vicinity of their insertion site (11%) or it evolved *de novo* from a cryptic intergenic promotor (86%). To assess expression of the newly inserted retroCNVs in the mouse populations, we used the transcriptomic dataset that was generated from the same individuals of the three natural *M. m. domesticus* populations from Germany, Massif Central of France and Iran (GE, FR_C and IR) for which also the genome sequences were obtained that we used for the retroCNV detection (Dataset S1B). To combine this information, we focused on the recently originated retrocopies present in the mm10 reference genome, as annotated in RetrogeneDB version 2 (27), since full length information for the inserted fragment is available for them. As the newly originated retrocopies are usually highly similar to their parental genes (25), we implemented an effective length (a proxy to the divergence to the parental gene) based approach to calculate their specific expression, by applying a high mismatch penalty strategy to distinguish the reads that can be perfectly and uniquely mapped to the new retrocopies (*SI Appendix*, Text S6).

Fifty-nine retrocopies with non-zero effective lengths across the three *M. m. domesticus* populations were included for this analysis. It should be noted that these retrocopies with non-zero effective lengths will be more diverged from the parental copy than those with zero effective length, but the expression levels of the latter ones cannot be quantified, since it is not possible to distinguish the reads that reliably map to the retroCNVs and those to the parental copy. We found that most of them (55 out of 59) are expressed in at least one tissue or at least one population (Dataset S4, summarized in *SI Appendix* Table S2). Most are expressed in multiple tissues, whereby the expression levels usually differ between the populations. This confirms the notion that the majority of retroCNV copies become transcribed after their insertion, although they responded differently to the regulatory context in their respective cell types and populations.

### RetroCNV effect on parental gene expression

Given that we have the expression data from the same animals for which we have the genome sequences, it was possible to ask whether the presence of a new retrocopy in a given individual would affect the expression of the parental gene in the same individual. To avoid any potential bias from population structure, we performed this line of analysis only for individuals from the same populations (FR_C and GE populations, separately). As these wild mice individuals were collected via a carefully designed sampling procedure, any possible effect from the genetic relatedness (or population substructure) among individuals should also be minimized (23, 24). We also restricted this analysis to the animals with singleton retroCNVs in each population, *i.e*., the cases where only one individual of a given population carried the retroCNV. This allowed us to use the remainder of the seven individuals from the same populations to calculate an average parental gene expression plus its variance, whereby all combinations of test versus reference individuals can occur. We used a Wilcoxon rank sum test to ask whether the presence of a retroCNV led to a significant expression change in the respective individual.

We firstly focused this analysis on the loss of expression, for which most likely antisense transcription of retroCNVs would silence the parental gene’s expression level (13). We found that 22% (GE) and 31% (FR_C) of the singleton retroCNVs have in at least one tissue a significant negative effect (FDR ≤ 0.05) on the expression of their parental gene (Table 1-Dataset S5A and S5B).

**Table 1:**
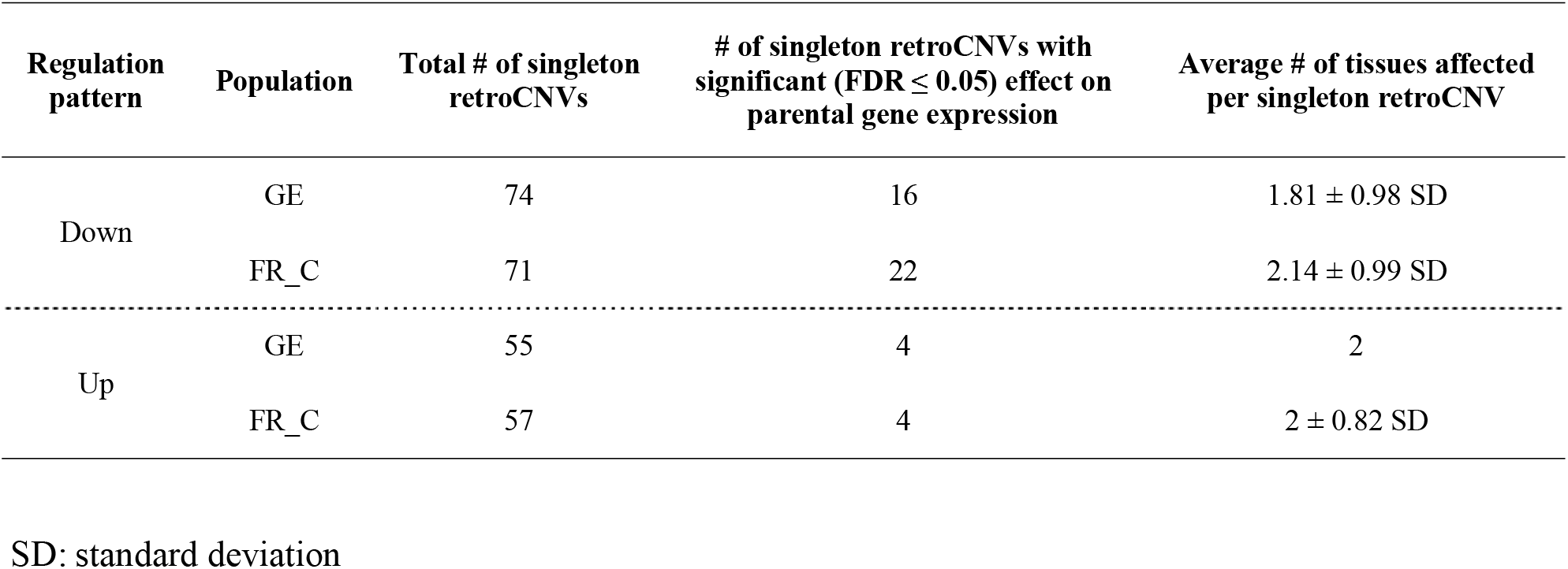
Singleton retroCNVs with significant effects on their parental genes’ expression in their population

Around three quarters of these retroCNVs (GE: 55/74; FR_C: 57/71) show truncated exons compared to their parental gene (Dataset S5C and S5D), and this allowed us to explore also upregulation effects on parental gene expression, since the expression level can be explicitly quantified based on the read fragments mapped to the exons that are unique to the parental gene. We found that about 7% of the singleton retroCNVs in both GE and FR_C populations have in at least one tissue a significant upregulation effect (FDR ≤ 0.05) on the expression of their parental gene (Table 1-Dataset S5C and S5D), hinting that retroCNVs could also functionally interfere with their parental gene expression through sponging regulatory microRNAs (15).

### Strand-specific expression of retroCNVs

To further assess whether the deleterious effects of retrocopies could be due to silencing effects from antisense transcribed copies, we generated a strand specific RNA-Seq dataset that allowed sense and antisense transcripts to be distinguished. For this we used five tissues from 10 males from the outbred stock of *M. m. domesticus* FR_C population. Note that these are different individuals than the ones used in Harr et al. (24), but from the same breeding stock of outbred animals. Hence, we could use the same reference genome set of retroCNVs (50 retroCNVs occurred in FR_C population), for which parental and retroCNV transcripts can be distinguished. We found that 42 of these 50 retroCNVs are transcribed in at least one tissue, but with an extreme bias towards sense-transcripts (Table 2 - Dataset S6). This applies not only to the number of transcribed retroCNVs per tissue, but also to the level of transcription (Dataset S6). Since only a low fraction (~3%) of retrocopies in mammals is expected to have inherited the promoter from the parental gene (33), it is unlikely that the direction of integration into the chromosomes could be biased to this extent. Hence, we interpreted this finding as a strong selection against retroCNV copies that showed antisense transcription, implying that they are affecting their parental genes via dsRNA silencing (13).

**Table 2:**
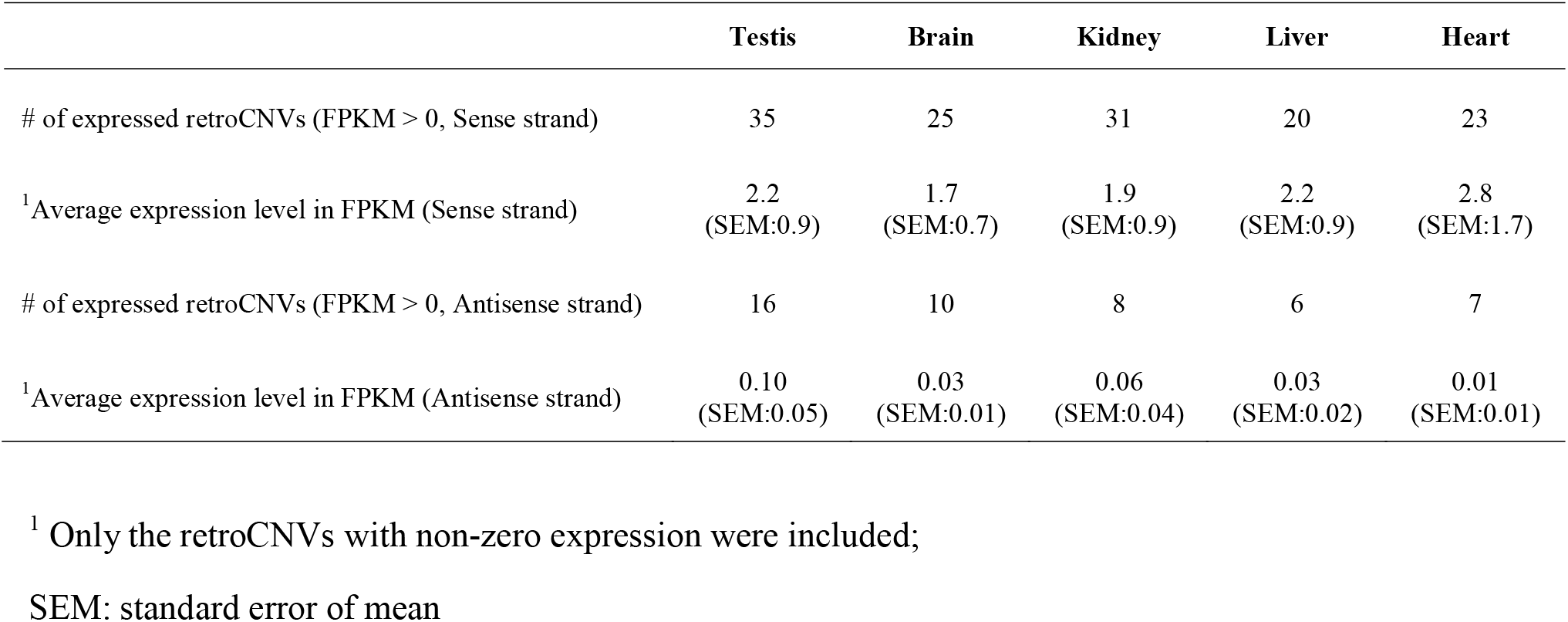
RetroCNV expression patterns in the strand-specific RNA-Seq dataset

|  | Testis | Brain | Kidney | Liver | Heart |
| --- | --- | --- | --- | --- | --- |
| # of expressed retroCNVs (FPKM > 0, Sense strand) | 35 | 25 | 31 | 20 | 23 |
| <sup>1</sup>Average expression level in FPKM (Sense strand) | 2.2<br>(SEM:0.9) | 1.7<br>(SEM:0.7) | 1.9<br>(SEM:0.9) | 2.2<br>(SEM:0.9) | 2.8<br>(SEM:1.7) |
| # of expressed retroCNVs (FPKM > 0, Antisense strand) | 16 | 10 | 8 | 6 | 7 |
| <sup>1</sup>Average expression level in FPKM (Antisense strand) | 0.10<br>(SEM:0.05) | 0.03<br>(SEM:0.01) | 0.06<br>(SEM:0.04) | 0.03<br>(SEM:0.02) | 0.01<br>(SEM:0.01) |

<sup>1</sup> Only the retroCNVs with non-zero expression were included;
SEM: standard error of mean

## Discussion

Our population based retroCNV analysis allowed a much deeper insight into the retrogene formation dynamics than was previously possible. Most importantly, we found that the primary origination rate of retroCNVs must be orders of magnitudes higher than previously assumed. At the same time, the data showed that many newly retroposed copies influence the expression of their parental genes and are mostly subject to negative selection, *i.e*., might be considered as “disease” alleles. Further, we showed that retroposed copies are not readily detected by previously established CNV detection procedures, *i.e*., their impact on generating deleterious mutations has been highly underestimated.

The comparison between very recently separated mouse populations provided the unique possibility to estimate primary retroposition rates, *i.e*., get an insight into the events that disappear over time from the populations due to negative selection. Such a disappearance of negatively selected variants is well known for functional SNPs and it has been shown to lead to a time dependence effect on measuring primary mutation rates. It was found that rates are much higher when very recent time horizons are studied, since the negative mutations can still segregate for some time in the populations (34). We have previously shown that this effect can also be traced in mitochondrial mutation patterns of mice after island colonization (35) and we observed it here for the comparisons of gene retroposition events between the most recently diverged populations.

Our rate estimates assume a more or less constant retroposition activity, rather than episodes of transpositions. As the main source of reverse transcriptase for retroposition, the LINE-1 elements are generally continuously active in mammals (2). In line with this, we observed similar numbers of retroCNV parental genes across all nine tested house mouse populations (Fig. 2B, median values range from 200 to 230), indicating a more or less constant rate of retroposition turnover activity. An episodic retroposition activity has been proposed to explain the relatively high birth rate of retrocopies in ancestral primates (36, 37), but in the light of our results, an alternative interpretation would be an enhanced retention rate of these retrocopies, possibly since some of them may have become involved in primate-specific adaptations.

Several of our analyses support the notion of strong negative selection acting on most new retroCNVs. In the comparison with the mutational spectra of SNP categories, we found that retroCNVs are even more deleterious than the category of the most deleterious SNPs, presumably because of their dominant effects. Most intriguingly, our data allowed directly to show the impact of new retroCNVs on the transcription of their parental genes. Especially the result on the strong transcriptional asymmetry bias among retroCNVs segregating in populations provides a direct clue why they may often be deleterious. Antisense RNA transcripts of retroCNVs would directly interfere with the function of the parental genes via RNA interference. While this may in a few cases have beneficial effects (13, 14) one can expect that it would mostly be deleterious. This would lead to a strong selection against highly expressed antisense retroCNVs and can thus explain why they are rare among segregating retroCNVs, or at least very poorly expressed. In our analysis of singleton retroCNV effects in populations (Table 1), we found between 22-31% having a negative effect on the expression of their parental genes. If one assumes that the primary integration of a retroCNV copy is random with respect to the orientation of transcription, half of the singletons could be in antisense direction, *i.e*., if the above percentage of negative effects is mostly due to antisense transcription, more than half of them are deleterious. Moreover, we have to assume that the most strongly deleterious ones are not represented in the sample, since they would be most quickly purged.

However, even sense copies could be deleterious due to dosage effects, or functional interference with their parental genes when truncated versions of the protein are produced, or through sponging regulatory micoRNAs (15). In our analysis of retroCNV effects in GE and FR_C populations (Table 1), we also found around 7% having a possibly negative effect caused by the upregulation of the expression of their parental genes. This is further supported by the observation that retroCNVs transcribed on the sense strand and antisense strand share the similar pattern of allele frequencies (*SI Appendix*, Fig. S12). Hence, while many previous reviews on retrogenes have focused of the evolutionary potential generated by retrogenes, the apparently strongly deleterious effects have been overlooked.

### Implications for human genetic disease studies

Our analysis thus suggests that the generation of retroCNV copies is a major contributor to the mutational load in natural populations. Mammalian genomes are estimated to carry up to about 1,000 deleterious SNP mutations per genome, mostly recessive ones (38). We found around 200 retroposition events per mouse genome, of which a substantial fraction is likely to have direct deleterious effects. This includes apparently most of the transcribed antisense copies, but also a fraction of the sense copies, given that we observe the strong purging of retroCNVs in comparison to SNPs. Most importantly, if the negative effects of retroCNVs are related to their transcription, only one allele would suffice to cause the effect, *i.e*., the negative effects are dominant. Accordingly, the retroCNV mutational load can be expected to be at least as large as that caused by (mostly recessive) SNPs.

A comparable retroCNV study in human populations (12) revealed also a very high rate of new retroCNVs, although about 3 times less (1663 retroCNV parental genes in house mouse populations versus 503 in human populations). However, the sequencing depth on the mouse samples is higher and our detection pipeline was further optimized. It is therefore reasonable to assume that the actual rate of retrocopy generation could be similar in humans and mice. Given their mostly dominant effect, this means that retrocopies may equally likely to cause a genetic disease than new SNP mutations. GWA studies of complex genetic diseases often find SNP associations in intergenic regions which are interpreted as regulatory variants. It is possible that such SNPs are in close linkage to an undetected retroCNV exerting a trans-regulatory influence on its parental gene and thus cause a disturbance of a genetic network. We note, however, that the variety of methods that are now available for SNP detection or structural variation detection do not yet include specific pipelines for retroCNV analysis (39). Although there are a few known cases where retroCNVs have caused a genetic disease through direct inactivation of genes (3, 40), a much more systematic approach to trace events caused by the transcriptional activity of retroCNVs seems warranted.

## Materials and Methods

### Glossary and definitions

Retroposed parental gene: A gene whose processed transcript is reverse-transcribed and re-inserted into the genome as a copy lacking the introns.

Retrocopy: A genomic DNA fragment that is generated from a parental gene through retroposition (RNA-based duplication).

RetroCNV parental gene: A retroposed parental gene for which the gene retroposition event is polymorphic in the target species.

RetroCNV (allele): The new retrocopy generated from a retroCNV parental gene which is polymorphic in the target species. On condition that the new retrocopy is present in the reference genome, it is denoted as the full length retrocopy; otherwise as the insertion site of the retrocopy in the genome when full length retrocopy is absent in the reference genome. In case that multiple retrocopies are generated from the same retroCNV parental gene, each retrocopy is considered as a distinct retroCNV.

### Reference genome sequences and gene annotation sources

We obtained the mouse reference genome sequence (mm10/GRCm38) and gene annotation data from Ensembl version 87 (41). This reference genome was built mainly based on the C57BL/6 mouse strain, which was derived from an inbred strain of the subspecies *Mus musculus domesticus*, but also included a small fraction of genome regions admixed from other house mouse subspecies, either due to inadvertent crosses in the initial breeding phase (42), or as remnants of natural introgression patterns (43).

We also retrieved the genome assembly sequence data for two out-group sister species (SPRET_EiJ_v1: *Mus spretus*; GCA_003336285.1: *Mus spicilegus*) from the NCBI Genbank database (44). Except *Mus spicilegus* (currently only at scaffold level), all other reference genomes were assembled at almost-complete chromosomal levels (45).

### Individual genomic sequencing datasets and short read alignment

We downloaded the whole-genome sequencing data from 96 wild individuals from 9 natural populations of 3 house mouse subspecies (listed in Dataset S1A) from either our previously generated dataset (24), or a publicly available dataset in the European Nucleotide Archive (ENA accession: PRJEB15873). In addition, we also included the whole genome sequencing data for 9 wild individuals from 2 out-group species for comparison (8, 24). Detailed description on these genomic sequencing data, including read length and sequencing coverage, can be found in Dataset S1A.

We aligned short sequencing reads from each individual genome to the mm10 reference genome sequence (Ensembl v87) in paired-end mode by using BWA mem (v0.7.15-r1140) (46), with default parameter settings, except the penalty for a mismatch (option “-B”) setting as 1, in order to compensate the sequence divergences of individuals from various populations and species. We only kept the alignment results to linear complete chromosomes. We further sorted alignment bam data by using the samtools sort function (v1.3.1) (47), and filtered PCR duplicates by using PICARD (v2.8.0) (http://broadinstitute.github.io/picard). The resulting alignment bam files were used for the further analysis.

### Identification of retroposed parental genes in the mouse inbred lines

We directly retrieved the datasets of retroCNV parental genes in the inbred strains of three house mouse subspecies (*Mus mus domesticus*: WSB_EiJ_v1; *Mus mus musculus*: PWK_PhJ_v1; *Mus mus castaneus*: CAST_EiJ_v1) provided in (18), which were generated based on short read genomic sequencing dataset by using a similar computational pipeline as shown below.

Additionally, we also identified recently retroposed parental genes found in the reference genome sequences of two out-group species, *i.e, Mus spicilegus* and *Mus spretus*. By refining previous strategies (25, 48), we applied a computational pipeline that searches mm10 reference genome exon-exon junction libraries (as shown below) against above out-group species reference genomes by using BLAT v36 with default parameters (≥90% alignment identity) (49), and only retained uniquely mapped regions. In case at least one exon-exon junction (≥30bp on each side) was uniquely mapped in the out-group species genome sequences, the corresponding gene was taken as a retroposed parental gene.

### Identification of house mouse specific retroCNV parental genes

Based on previous approaches (18, 19, 25), we developed a refined computational pipeline for the discovery of retroCNV parental genes based on the short read sequencing datasets from individual genomes (*SI Appendix*, Fig. S1). This pipeline combines both exon-exon and exon-intron-exon junction read mapping strategies to identify gene retroposition events, and the discovery process is independent of the presence of newly generated retrocopies in the reference genome. A more detailed description on the discovery of retroCNV parental genes can be found in *SI Appendix* Text S3.

In this study, we only focused on house mouse specific retroCNV parental genes, which were defined as retroposed parental genes detected in at least one house mouse individual, while absent in both the outgroup species (*Mus spicilegus* and *Mus spretus*), neither in wild individual genome sequencing data nor in the inbred reference genomes.

### Detection of retroCNV alleles

Based on the above detected house mouse specific retroCNV parental genes, we performed detection of retroCNV alleles at individual genome level. The presence statuses of retrocopies those are annotated in the mm10 reference genome and the insertion sites for those retrocopies absent in the reference genome were analyzed separately (*SI Appendix* Text S4).

For retroCNVs of which the insertion sites inferred from different individuals that were from the same retroCNV parental gene, we used 1kb as the clustering distance threshold (also required on the same chromosomal strand) to define the same gene retroposition events. Accordingly, the multiple retrocopies (with distinct insertion sites) from the same retroCNV parental gene were taken as distinct retroCNVs. In total, we included 2,025 unique house mouse specific retroCNVs (*SI Appendix*, Fig. S6) for the further analysis.

Additionally, we also checked for the positive evidence of retroCNV absence (*i.e*., alignments spanning the retroCNV alleles) for the occasions where the retroCNV alleles (*i.e*., positive evidence of retroCNV presence) cannot be detected for some individuals. We searched for proper paired-end alignments (with both correct orientation and expected mapping distance) that are uniquely mapped (same criteria to define unique alignment as above), and spanning the estimated insertion site (retroCNV absent in the mm10 reference gnome) or both sides of the flanking regions (retroCNV annotated in the mm10 reference genome) of retroCNV allele. Similarly, we required at least two supporting alignments to call the positive evidence of retroCNV absence. For around 95.5% of the cases where the retroCNV alleles cannot be detected, we can observe the positive evidence of retroCNV absence, confirming the reliability of the above retroCNV detection pipeline. Consequently, we refined a new set of 1,551 retroCNVs (Dataset S3), including only the retroCNVs that show both positive evidence of retroCNV presence and positive evidence of retroCNV absence in all 96 tested house mice individuals.

### Comparison of the allele frequency pattern between retroCNVs and SNPs

We followed the general GATK version 3 Best Practices (50) to call SNP variants (*SI Appendix*, Text S5), and only kept the SNP variants with unambiguous ancestral states in out-group species (*i.e*., same homozygous genotype for all 9 tested individuals from 2 out-group species), while with alternative allele in house mouse individuals for further analysis. We predicted the functional effects of each SNP by using Ensembl VEP v98.2 (32), based on the gene annotation data from Ensembl version 87 (41). Consistent with Ensembl variation annotation (41), we categorized these SNPs into four groups given their predicted impacts: 1) High effect - SNPs causing the gain/loss of start/stop codon or change of the splicing acceptor/donor sites; 2) Moderate effect - SNPs resulting in a different amino acid sequence; 3) Low effect - SNPs occurring within the region of the splice site, changing the final codon of an incompletely annotated transcript, changing the bases of start/stop codon (while start/terminator remains), or where there is no resulting change to the encoded amino acid; 4) Modifier effect - SNPs occurring within the genes’ non-coding regions (including UTR, intron, up/downstream), or intergenic regions.

For both retroCNV and SNP alleles, we calculated their allele frequency at three difference levels (subspecies, population and individual) by counting individuals with positive evidence of each allele, without distinguishing the homozygous and heterozygous genotype status. We performed two independent analyses on both sets of retroCNVs as defined in the above section, *i.e*., the overall 2,025 retroCNV dataset and the other refined subset of 1,551 retroCNVs, and showed the same allele frequency pattern (Fig. 3 and *SI Appendix*, Fig. S11). The former dataset was then taken as representative for all the following analysis.

The frequency spectrum of house mouse retroCNVs was further compared with the site frequency spectrum of SNPs from the four above defined categories. We quantified the distances between spectrum distributions by using two sided Kolmogorov-Smirnov tests, and calculated the statistical significances of the fraction of individual private variants by using Fisher’s exact tests.

### Transcriptional profiling of retroCNVs

We used two different sets of transcriptomic sequencing data for the transcriptional profiling of retroCNVs: 1) one non-strand-specific RNA-Seq dataset from our previously published data (24); 2) one strand-specific RNA-Seq dataset newly generated in the present study. The detailed description about these two datasets can be found in Dataset S1B and S1C.

In order to accurately quantify expression levels, we focused on the recent retrocopies present in the mm10 reference genome (originated from house mouse specific retroCNV parental genes, and with sequence identity ≥ 95% compared with their parental genes), for which the information was directly inferred from RetrogeneDB version 2 (27). As the recently originated retrocopies are usually highly similar to their parental genes, we implemented an effective-length based approach to calculate their expression values (25). We calculated the effective length of each retrocopy as the number of uniquely mapping locations in the retrocopy region, and only kept 59 retrocopies (50 of them are present in FR_C population, thus were used for strand-specific RNA-Seq dataset analysis) with non-zero effective length for further analysis. The normalized FPKM value for each retrocopy within each tissue was calculated on the basis of the above computed effective length of each retrocopy. A detailed description on this line of analysis is provided in *SI Appendix* Text S6.

### The impact on parental gene expression from singleton retroCNVs

Given the limit number of mice individuals (N=4) with matched genomic/transcriptomic sequencing dataset in Iran population (Dataset S1B), we restricted this analysis to the animals with singleton retroCNVs in the FR_C (71 singleton retroCNV parental genes) and GE (74 singleton retroCNV parental genes) populations (N=8 for both populations) only, *i.e*, the cases where only one individual of a given population carried the retroCNVs. In case that one singleton retroCNV parental gene has multiple retroCNV alleles (*i.e*, ≥2 detected insertion sites), the effects of all these alleles were combined (Dataset S5), since it is unlikely to separate their own effects. Moreover, the singleton retroCNV parental genes that have no detectable insertion site (likely landing in the repetitive genomic region) were also included for analysis here.

As shown in *SI Appendix* Text S7, the analyses on down- and up-regulation impact on parental gene expression were performed separately. For both down- and up-regulatory impact analysis, we used Wilcoxon rank sum test to calculate the significance (p-value) of the parental gene expression change between singleton retroCNV carrier and non-carriers, and the significant expression changes after multiple testing corrections (FDR ≤ 0.05) were also included.

### Data access

The raw strand-specific RNA-Seq data generated in this study have been submitted to the European Nucleotide Archive (ENA; https://www.ebi.ac.uk/ena) under study accession number PRJEB36991.

## Acknowledgements

We appreciate Peter Keightley and Guy Reeves for reading through the manuscript and providing helpful comments. We are grateful to Julien Dutheil for valuable suggestion on statistical analysis. We thank Yuanxiao Gao for valuable suggestions on data presentation and visualization. We appreciate the laboratory members for helpful discussions and suggestions. Computing was supported by the Wallace HPC cluster of the Max-Planck Institute for Evolutionary Biology. This work was supported by institutional funding through the Max Planck Society to DT.

## Supplementary Information Text

### Text S1 Parameter optimization for the computational discovery of retroposed parental genes

The genomic sequencing datasets from wild mouse individuals varied in read length, sequencing coverage and sequence divergence to the mm10 reference genome (*SI Appendix*, Table S1 and Dataset S1A). These distinct features needed to be considered for the parameter optimization of the discovery of retroposed parental genes. Several parameters were included in the discovery pipeline: 1) criteria to define unique read mapping; 2) spanning read length (both exon and intron side) to define reliable supporting evidence; 3) number of supporting evidences to define “true” gene retroposition event.

In order to detect recent retroposed parental genes in each individual genome, only mapped reads with alignment identity ≥95% and mapping quality MapQ ≥20 were included, and the alignment identity parameter further adjusted based on individual genome sequence divergence to mm10 reference genome (see below text). Furthermore, a customized Perl script was implemented to compare the alignment scores of multiple-hit reads and retained only the uniquely-mapping reads if the difference between the best and second best alignment score is ≥5 (1).

To determine the optimal spanning read length, we searched the built exon-exon junction database against mm10 reference genome by using BLAT v36 with default parameters (2), and only kept the returning searching hits with ≥95% alignment identity. Furthermore, we selected alignment results for those spanning at least 10bp, 20bp, 30bp, 40bp and 50bp on both sides of consecutive exons. The corresponding genes of these aligned exon-exon junctions were putative retroposed parental genes. In case that one retroposed parental gene was not included in any of those three published resources: RetrogeneDB (v2) (3), UCSC (RetroV6) (4) and GENCODE (v20) (5), it was considered as a false positive prediction. As shown in *SI Appendix* Fig. S2, the false discovery rate (FDR) stabilized at spanning size of 30bp (~ 2.5%), with no major reduction of FDR as the spanning size increased. Therefore, this size of 30bp was set as the basis of threshold selection of spanning read length for both exon and intron side. The relative longer read length of our sequencing datasets permitted spanning size threshold that was much larger than previous reports (15bp in (1), 5bp in (6)), and this helped to define a more reliable retroposed parental gene dataset. This parameter was further adjusted according to the features of individual sequencing datasets (see text below).

In order to assess the potential influence of sequencing dataset features on the discovery of retroposed parental genes, we conducted a systematic simulation analysis by incorporating various aspects of sequencing datasets: 1) sequence divergence to mm10 reference genome; 2) sequencing read length; 3) sequencing depth (*SI Appendix*, Fig. S3). We randomly selected 1000 consecutive exon-exon junction sequences (corresponding to 1000 “putative” retroposed parental genes) and then randomly inserted them into mm10 reference genome sequences. On the basis of real individual sequencing data (*SI Appendix*, Table S1), four types of random mutation rate (u=0.00, 0.01, 0.02, 0.03) at single nucleotide sites, two types of sequencing read length (75bp/100bp) and three types of sequencing depth (10X, 20X and 30X) were simulated to generate in total 24 distinct sequencing datasets. The simulation of genomic sequencing datasets was performed by using Art_illumina (v 2.5.8) (7), with pair-end read simulation mode (Insert size for 75 bp reads: 150bp; Insert size for 100 bp reads: 200bp). The above computational pipeline (*SI Appendix*, Fig. S1) was applied to identify retroposed parental genes from these 24 simulated genomic sequencing datasets.

As expected, in case that uniform setting of parameters (Alignment identity ≥95%; Spanning read length ≥30bp; Number of supporting evidences ≥3) were applied, genomic sequencing datasets with longer read length and higher sequencing depth tended to have larger recall rate (sensitivity) (*SI Appendix*, Fig. S4), while mutation rates (in relation to sequence divergence) only played minor roles after increasing to 0.03. Therefore, the discovery parameters were adjusted with the rule of lowering strictness for datasets with shorter read length, larger mutation rate and lower sequencing depth (*SI Appendix*, Fig. S3). After the optimization of parameters, relative constant recall rates (≥95%) were observed for all 24 simulated sequencing datasets (*SI Appendix*, Fig. S5). Accordingly, the detection parameters for the real sequencing datasets were also adjusted based on the same rules (Dataset S1A). It was less likely that the adjustment of detection parameters could introduce more false positive discoveries, with the fact that most real genomic sequencing datasets have either relatively long read length or high sequencing coverage (Dataset S1A).

### Text S2 Possible technical issues on the detection of retroCNVs

Since the detection of retrocpies relies on the identification of pieces of retroposed genes (*i.e*., exon-exon junction events), it is likely that other structural mutation mechanisms might introduce bias in the discovery of retrocpoies. In this section, we explore in detail the possible situations that may cause false discovery of retrocopies, and to what extent they might affect our conclusions.

*SI Appendix* Fig. S7 illustrates the possible scenarios to introduce false discovery of retrocopies. The first scenario refers to the possibility the structural differences (*i.e*., intron deletion) of the gene alleles could be taken as gene retroposition (*SI Appendix*, Fig. S7A). Firstly, we need to clarify that our retroposition event detection pipeline requires both the presence of intron loss events and presence of the parental gene allele (*SI Appendix*, Fig. S1). Therefore, the only possibility is for the heterozygous allele of intron deletion. Secondly, it should be considered how less likely other structural mutation mechanisms (instead of gene retroposition) could cause the intron deletion at the nearly exact boundary of exon-intron and intron-exon, with less than 3bp differences (otherwise will not be detected by our pipeline). Indeed, our simulation analysis has shown that the false discovery rate of intron loss event (*i.e*., presence of intron loss, but not from gene retroposition) is <3% for a single pair of consecutive exons (*SI Appendix*, Fig. S2). We also found that majority (>71%) of detected gene retroposition events involve >=2 intron loss events (*i.e*., corresponding to multiple pairs of consecutive exons), and therefore the possibility for the structural intron deletions for these gene retroposition events would be even much lower (<0.09%). Lastly, no retroCNV allele will be called for intron deletion events (as no discordant alignments can be found to infer new insertion site), since the structural deletion will not change the genomic coordinate of the gene locus. On the basis of these arguments, we rule out the possibility that the structural differences of the same gene alleles can largely affect our finding in the present study.

The second scenario refers to the possibility that multiple retroCNV alleles could be called due to the DNA duplication of the existing “real” retrocopy (*SI Appendix*, Fig. S7B). Firstly, it is worth noting that this argument only potentially affect a low fraction (<4%) of the retroposition event calling, as it is reported that only 3.9% of retroCNV parental genes show more than one retroCNV allele in our study (*SI Appendix*, Fig. S8). Secondly, it can be argued that the possibility of the recurring structural mutations (gene retroposition and DNA duplication) at the same genomic region should be low, especially in this rather short evolutionary time scale (<0.5 MYA). For instance, only around 9.8% of annotated retrocopies in mm10 reference genome from RetrogeneDB v2 (8) are found to overlap the DNA segmental duplication region annotated in UCSC database. Lastly, the DNA segmental duplication usually involve large DNA segments, thus the flanking regions of the retrocopy are also highly likely to be included in the rearrangement event (*i.e*., copy to a new genomic location). Consequently, we would not call such cases as separate events, they would only show up as higher read coverage, but we have not included this as a detection criterion. Hence, we conclude that the possibility of a secondary DNA duplication of the newly inserted retrocopy is rather low, and not likely to heavily change our findings.

The last refers to the possibility that there could be a secondary break (*i.e*., inversion or DNA segmental translocation) in a recently transposed retrocopy (*SI Appendix*, Fig. S7C). In this case the two halves could be counted as two events, rather than one. First of all, it should be noted that they would still identify the same corresponding parental gene. Secondly, similar to the first scenario, this scenario would also only relate to a low fraction (<4%) of called retroposition events, and additionally the possibility of the recurring structural mutations should be rather low as well. Thus, we could also rule out the possibility that a secondary break in a recently transposed retrocopy can largely affect our conclusions.

Overall, other structural mutation mechanisms are either less likely, or could only introduce a very low fraction of false discovery of retrocopies, compared with the large number of retrocopies detected at genome-wide scale. Therefore, we conclude that our main conclusions in the present report should not be substantially influenced by a couple of false discovery of retrocopies (if any), due to other structural mutation mechanisms.

### Text S3 Identification of house mouse specific retroCNV parental genes

On the basis of previous approaches (1, 9, 10), we developed a refined computational pipeline for the discovery of retroCNV parental genes based on the short read sequencing datasets from individual genomes (*SI Appendix*, Fig. S1). This pipeline combines both exon-exon and exon-intron-exon junction read mapping strategies to identify gene retroposition events, and the discovery process is independent of the presence of newly generated retrocopies in the reference genome. The reliability of this approach has been experimentally verified with PCR techniques in a couple of previous studies (1, 9).

We firstly generated an exon-exon junction database, by extracting all possible consecutive exon-exon junction sequences (100bp from each side, or shorter for small exons, but ≥50bp) for all 20,378 protein coding genes with ≥2 exons annotated in the mm10 reference genome based on Ensembl v87 (11). Similar to the mapping procedure to the reference genome, all the short sequencing reads from each individual genome were aligned to the exon-exon junction database by using BWA mem (v0.7.15-r1140) (12), with default parameter settings except the penalty for a mismatch (option “-B”) setting as 1. Given the rather short length (≤ 200bp) of exon-exon junction sequences, the alignment to the exon-exon database was done in single-end mode. We defined uniquely mapped reads (for both exon-exon junction dataset mapping and reference genome mapping) as the ones satisfying both of the two criteria: 1) mapping quality ≥20; 2) the difference between the best and second best alignment scores ≥5 (1).

A retroposed parental gene is called, in case that both the intron loss events (*i.e*., exon-exon junction mapping) and the presence of a parental gene (*i.e*., exon-intron and intron-exon junction mapping) can be observed in the same individual sequencing dataset (1). On the basis of our simulation results in *SI Appendix* Text S1, for the calling of both events, we required at least 2 distinct supporting reads (to be adjusted according to individual genomic sequencing dataset as shown below) from the same individual genome, that span at least 30bp on each side of exon-exon/exon-intron-exon junction.

Given the distinct features (*i.e*., read length, sequencing coverage, and divergence to the mm10 reference genome) of the short read sequencing dataset for the individuals from each population (*SI Appendix*, Table S1), we further conducted a comprehensive simulation analysis to tailor the discovery pipeline for each population individuals with different parameter settings (*SI Appendix*, Text S1): 1) alignment identity; 2) minimum spanning length on each side of junction; 3) minimum number of supporting reads. The optimized parameter setting for the discovery of retroposed parental genes in each individual genome is provided in Dataset S1A.

### Text S4 Detection of retroCNV alleles

Based on the above detected house mouse specific retroCNV parental genes, we performed detection of retroCNV alleles at individual genome level. The presence statuses of retrocopies those are included in the mm10 reference genome and the insertion sites for those ones absent in the reference genome were analyzed separately.

For the retrocopies present in the mm10 reference genome (with sequence identity >=95% to their parental gene), we used the ones annotated in RetrogeneDB version 2 (8). We searched for proper paired-end alignments (with both correct orientation and expected mapping distance) from each house mouse individual sequencing dataset, that have one read uniquely mapped to the flanking region of the annotated retrocopy and the other read mapped within the focal retrocopy region (unique mapping not required). Two criteria were required to define the unique alignments to the flanking region: 1) mapping quality ≥20; 2) the difference between the best and second-best alignment scores ≥5. A presence of retrocopy is called, if there are at least two supporting reads on both flanking sides or at least four supporting reads on either flanking side.

In order to detect the insertion sites for the retroCNVs that are absent in the mm10 reference genome (*i. e*., full length retrocopy is not available), we applied a discordant alignment based approach by using the above paired-end read alignments from each individual genome sequencing data (13, 14). We searched for paired-end alignments in proper orientation, and with one read uniquely mapped (*i.e*., anchor read) within exonic sequences of the parental gene and the other read uniquely mapped to a distinct genomic region, *i.e*., on a different chromosome or on the same chromosome but with an unexpected mapping distance (> 2 x average insert size of the paired-end library). Similarly, a unique alignment was required to meet both criteria: 1) mapping quality ≥20; 2) the difference between the best and second-best alignment score ≥5. The procedure of the clustering of the above discordant alignments was performed by following (13, 14), with 500 bp as the cut-off for average linkage distance to stop clustering. The insertion site was taken as the middle point of the cluster. An insertion site would be considered to be valid, if there are at least two supporting reads on both sides (*i.e*., strands), or at least four supporting reads on either side (14).

### Text S5 Single nucleotide polymorphism (SNP) calling

We followed the general GATK version 3 Best Practices (15) to call variants. Specifically, we realigned the above PCR-duplicates filtered alignment bam files around the indels (with flag “PASS”) detected from the Mouse Genome Consortium (v5) (16) with GATK (v3.7), and recalibrated base quality scores with by using SNP variants (with flag “PASS”) founded in the Mouse Genome Consortium (version 5) (16) to get analysis-ready reads.

In the first, we called raw genetic variants for each individual using the HaplotypeCaller function in GATK (v3.7), and then jointly genotyped genetic variants for all the individuals using GenotypeGVCFs function in GATK (v3.7). We only retained bi-allelic SNP variants that passed the hard filter “QD < 2.0 || FS > 60.0 || MQ < 40.0 || MQRankSum < −12.5 || ReadPosRankSum < −8.0 || SOR > 3.0”, and dropped the variants with missing calling value in any individual. We only kept the SNP variants with unambiguous ancestral states in out-group species (*i.e*., same homozygous genotype for all 9 tested individuals from 2 out-group species), while with alternative allele in house mouse individuals for further analysis.

### Text S6 Transcriptional profiling of retroCNVs

We used two different sets of transcriptomic sequencing data for the transcriptional profiling of retroCNVs: 1) one non-strand-specific RNA-Seq dataset from our previously published data (17); 2) one strand-specific RNA-Seq dataset newly generated in the present study. The detailed description about these two datasets can be found in Dataset S1B and S1C.

The non-strand-specific RNA-Seq dataset used the same individuals which are included in the above whole genome sequencing data of *M. m. domesticus* (17). Four individuals in Iran population were excluded for analysis due to imperfect match between genomic/transcriptomic datasets. We downloaded the transcriptomic sequencing data of 10 tissue samples (*i.e*., Brain, Gut, Heart, Kidney, Liver, Lung, Muscle, Testis, Spleen, Thyroid) from 20 mice individuals of 3 natural populations (Germany population (GE): 8 individuals; France Massif Central population (FR_C): 8 individuals; Iran population (IR): 4 individuals). With a few exceptions, most of these individuals have expression profiling data from all these 10 tissues (Dataset S1B).

To study the orientation-dependent transcription of the retroCNVs, we generated a strand-specific RNA-Seq dataset from 10 male individuals of the France Massif Central population (FR_C). The (whole) brains, hearts, livers (right medial lobes), (right) kidneys, and (right) testes from ten 24-26 weeks old males were carefully collected and immediately frozen with liquid nitrogen. Total RNAs were purified using RNeasy 96 Universal Tissue Kit (Catalog no. 74881), and sent to Competence Centre for Genomic Analysis in Kiel for stranded mRNA library preparation and sequencing on Illumina NovaSeq 6000 (2 × 150bp). For both datasets, fastq files were trimmed with Trimmomatic (0.38) (18), and only paired-end reads passed filtering process were used for further analyses.

In order to accurately quantify expression levels, we focused on the recent retrocopies present in the mm10 reference genome (originated from house mouse specific retroCNV parental genes, and with sequence identity ≥ 95% compared with their parental genes), for which the information was directly inferred from RetrogeneDB version 2 (8). As the recently originated retrocopies are usually highly similar to their parental genes, we implemented an effectivelength based approach to calculate their expression values (1). Firstly, we simulated 5000X 100bp paired-end sequencing reads on the basis of retrocopies’ sequences (including 500bp up/downstream flanking regions) with Art_illumina (v 2.5.8) (7), and then re-mapped them to the combined reference sequences (mm10 reference genome + transcript sequences of the parental genes of these retrocopies) using BWA mem (v0.7.15-r1140). In order to distinguish from parental genes, we applied a high penalty for the mismatch (or insertion/deletion, −B 10, - O [10,10]) for the mapping process. Only the uniquely aligned reads (a. mapping quality ≥ 20; b. perfectly match in the retrocopy regions, but at least one site mismatch for the second-best alignment to the parental gene, AS-XS ≥ 11) were included. We calculated the effective length of each retrocopy as the number of uniquely mapping locations in the retrocopy region, and only kept 59 retrocopies (50 of them are present in FR_C population, thus were used for strand-specific RNA-Seq dataset analysis) with non-zero effective length for further analysis. Secondly, we pooled the transcriptomic sequencing data of the retroCNV carriers of the same population individuals for each tissue, and mapped them to the same combined reference sequences with the same BWA mem pipeline as mentioned above. For the strand-specific RNA-Seq dataset, we counted the reads of all those 10 tested individuals from the two strands (sense and antisense strands relative to their parental genes) separately. The normalized FPKM value for each retrocopy from each strand within each tissue was calculated on the basis of the above computed effective length of each retrocopy.

### Text S7 The impact on parental gene expression from singleton retroCNVs

To explore the negative impact on parental gene expression, we compared the expression level of the focal parental gene in the mice individual with the singleton retroCNV with those in the reminder of seven retroCNV non-carriers from the same population. Firstly, we mapped the non-strand-specific RNA-Seq reads from each individual/tissue from FR_C and GE population individuals to mm10/GRCm38 reference genome sequence with HISAT2 (2.1.0) (19), taking advantage of the mouse gene annotation in Ensembl v87 by using the --ss and --exon options of the hisat2-build. Then we counted the fragments mapped to the annotated genes with featureCounts (1.6.3) (20). Finally, we calculated the expression level in FPKM (Fragments Per Kilobase of transcript per Million mapped reads) for each annotated coding gene within each individual/tissue on the basis of gene’s transcript length and the sequencing dataset size from each individual/tissue.

Based on the information of exon-exon junction and the estimated insertion site of retroCNV alleles, we found that around three quarters of the singleton retroCNVs (GE: 55; FR_C: 57) harbor at least one truncated exon (Dataset S5), compared with their respective parental gene. The existence of exon(s) unique to retroCNV parental genes allows to explore the retroCNV allele’s up-regulatory impacts on parental gene expression, as we could use this information to explicitly quantify the expression level of retroCNV parental gene. Based on the HISAT2 alignment bam data from above, we counted the fragments that mapped to the annotated exons with featureCounts (1.6.3) (20). We calculated the expression level of each retroCNV parental gene in FPM (Fragments per Million mapped reads) within each individual/tissue on the basis of the fragment counts mapped to the unique exon(s) to the parental gene (relative to retroCNV allele) and the mapped sequencing dataset size from each individual/tissue. In case of multiple transcripts associated with one parental gene, the longest one was taken as representative.

### Text S8 Genic copy number variation (CNV) calling

We predicted CNV calls for each individual based on the above PCR-duplicates filtered alignment bam files by using the sequencing read depth based approach implemented in the program CNVnator v0.3.3 (21), as the reliability of this approach has been experimentally confirmed (22). We chose the optimal bin size for each individual, such that the ratio of the average read-depth signal to its standard deviation was between 4 and 5 (21). Bin size ranged from 100-1,500bp and was inversely proportional to genome coverage. Following the convention in (22), we discarded the CNV calls below 1kb in length and intersecting annotated gaps in the reference genome, and defined “genic CNVs” as the genes for which at least one whole transcription unit is completely contained within CNV calls. Similarly, house mouse specific “genic CNVs” were defined as the ones (duplications or deletions) can be found in at least one house mouse individuals, but not in any individual from outgroup species. We computed the fractions of overlapping retroCNV parental genes and genic CNVs for each house mouse individual from nine natural populations separately.

**Fig. S1.**
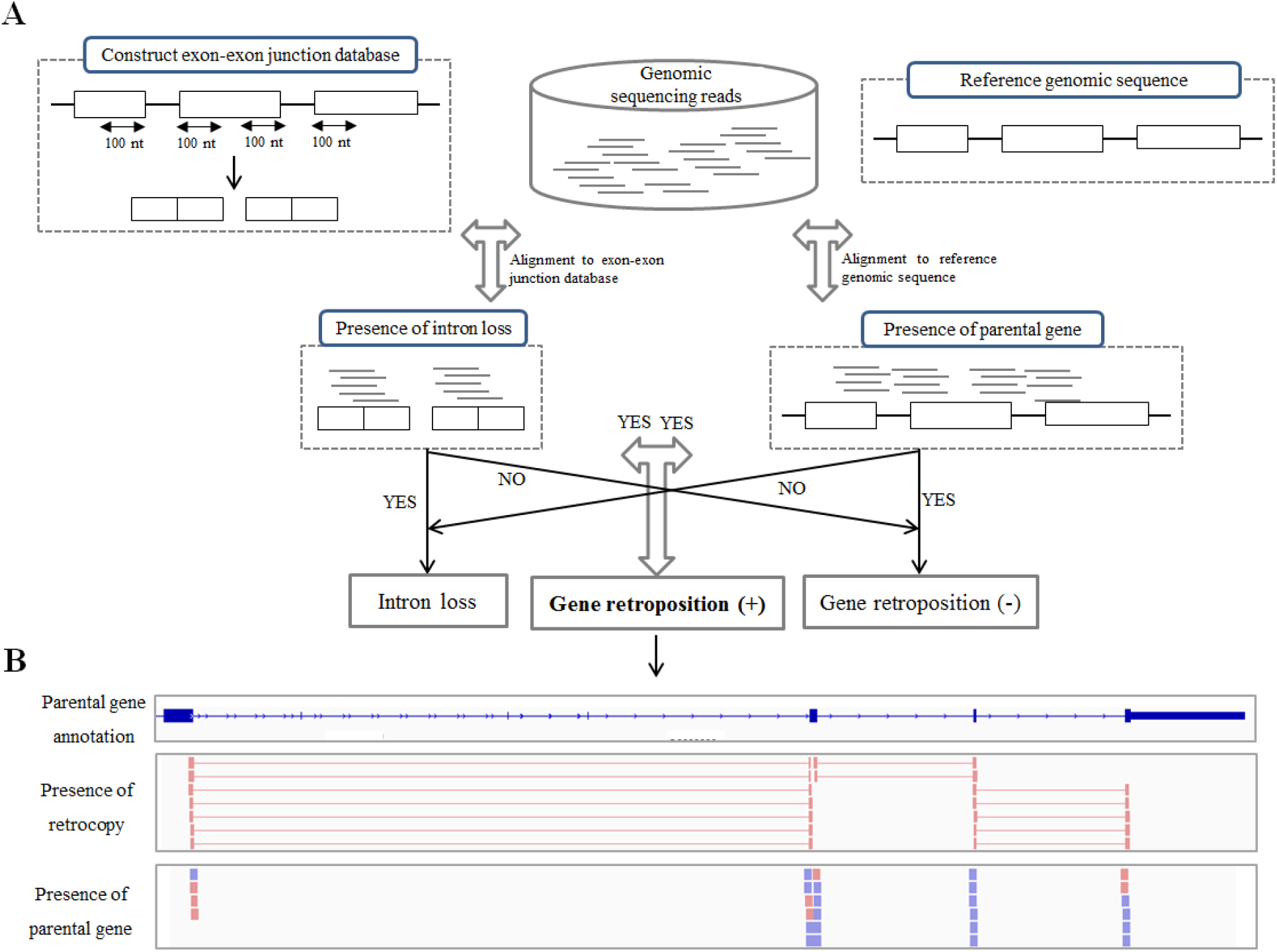
Computational pipeline for the discovery of retroposed parental genes. (A) shows a simplified flow chart of our computational calling pipeline. (B) shows an example of gene retroposition event in the view of an IGV snapshot.

**Fig. S2.**
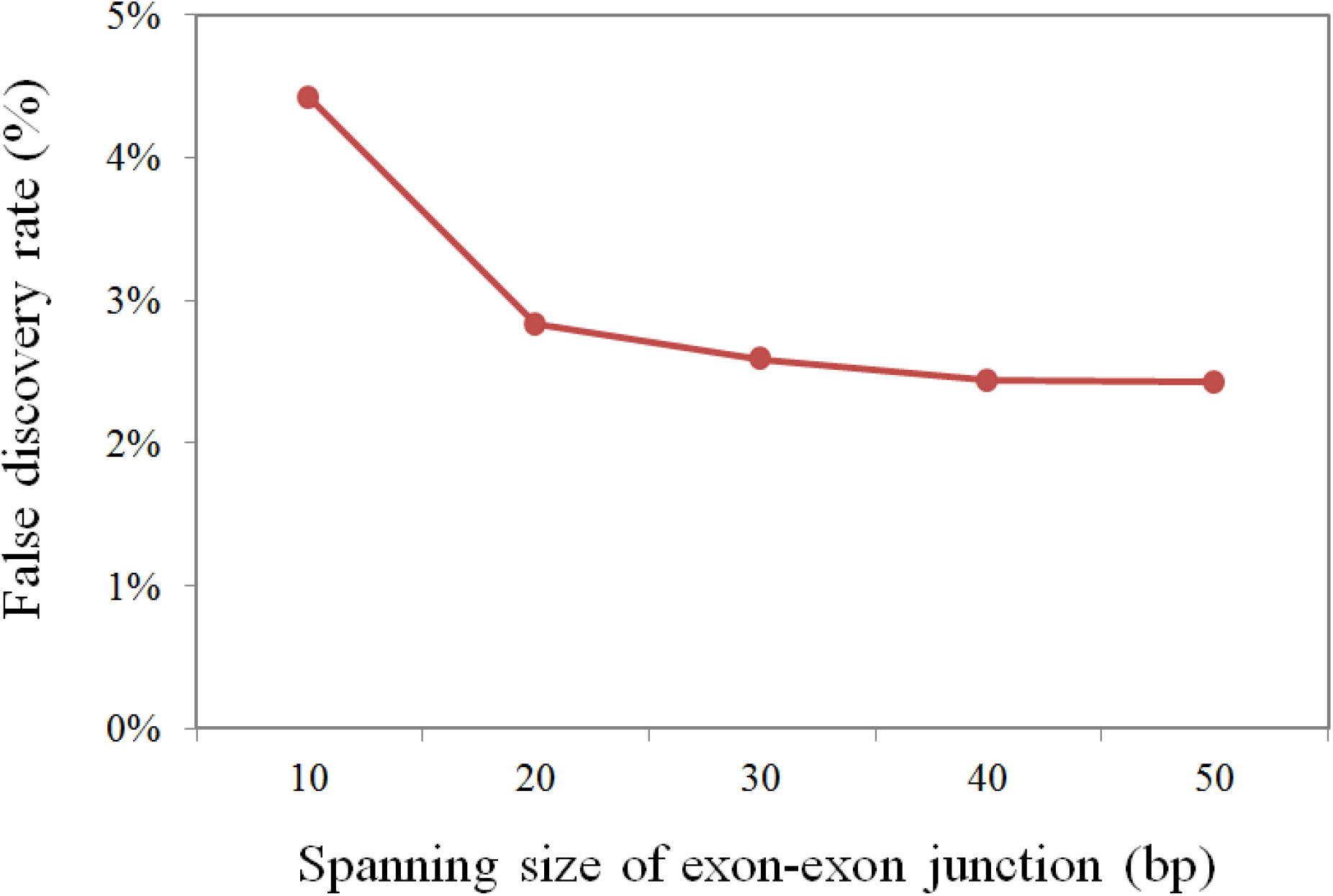
False positive discovery rates of retroposed parental genes based on the different spanning sizes cutoffs. False discovery rate was calculated as the number of predicted retroposed genes that were not included in any of those three previous annotation datasets (RetrogeneDB v2, UCSC Retro V6, and GENCODE v20), divided by the total number of predicted retroposed parental genes.

**Fig. S3.**
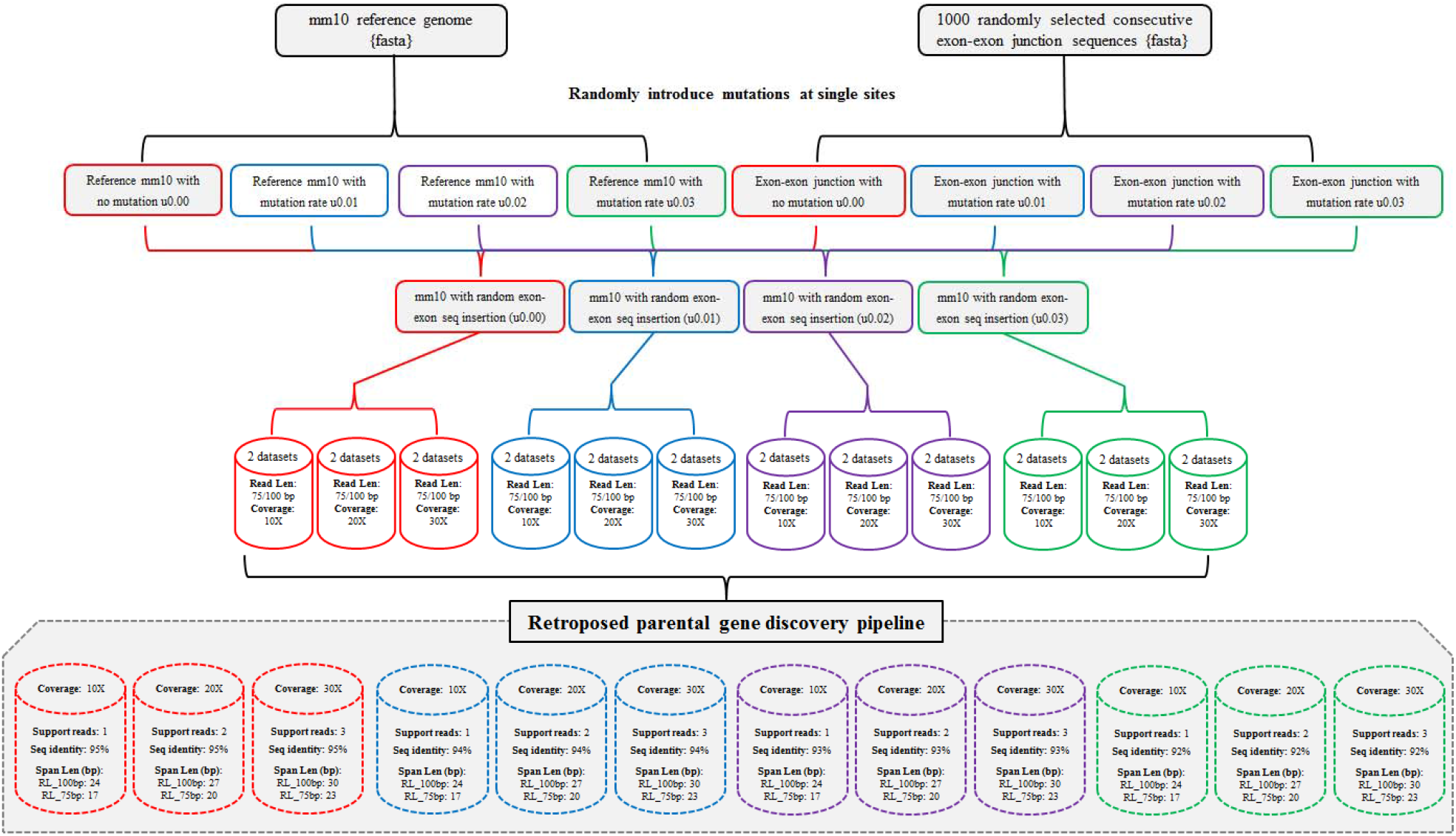
A flow chart to show the parameter optimization of the retroposed parental gene detection pipeline. In total, 24 genomic read sequencing datasets were simulated on the basis of the features of real individual sequencing datasets. The bottom panel shows the optimized parameters for the discovery of retroposed parental genes for different genomic sequencing datasets.

**Fig. S4.**
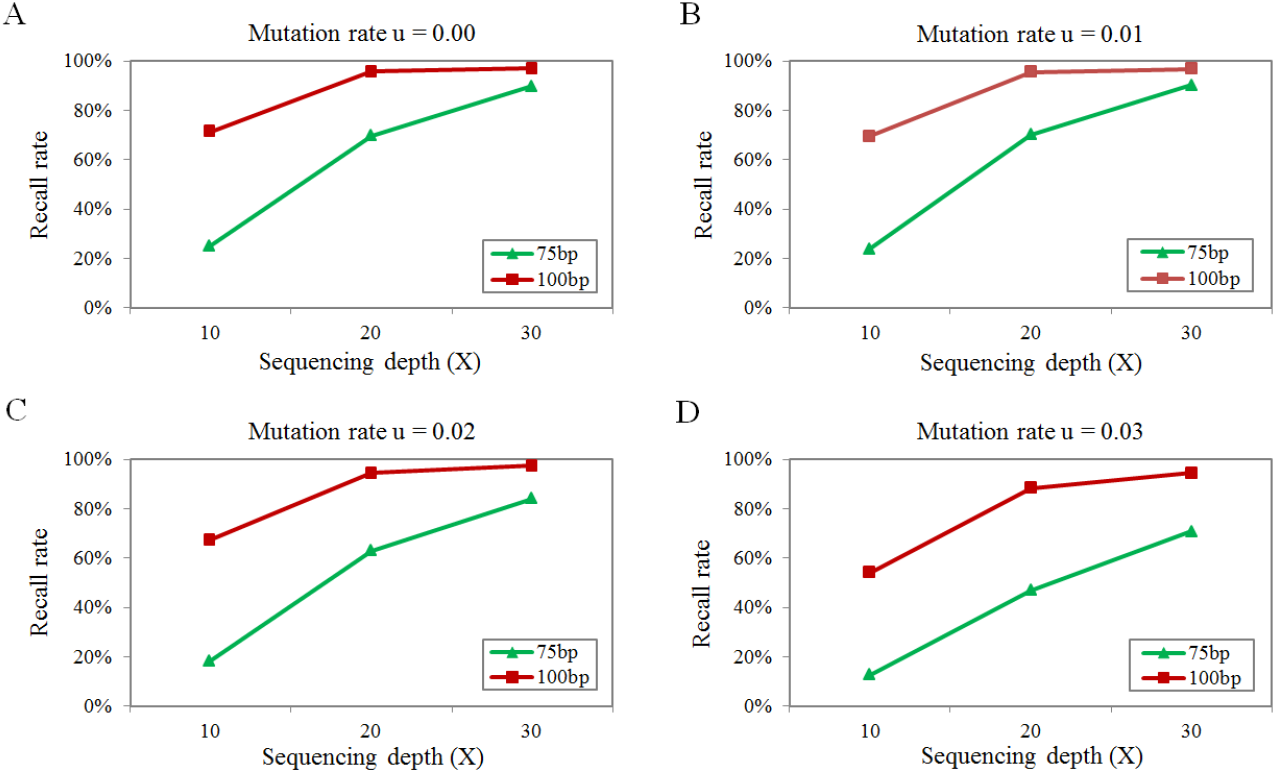
Recall rates of the discovery of retroposed parental genes with uniform setting of parameters. For all simulated genomic sequencing datasets, the same settings of parameters were applied: Alignment identity ≥95%; Spanning read length ≥30bp; Number of supporting evidences ≥3. The recall rate for each simulated genomic sequencing dataset was calculated as the fraction of identified ‘true’ retroposed parental genes discovered from 1,000 random simulations.

**Fig. S5.**
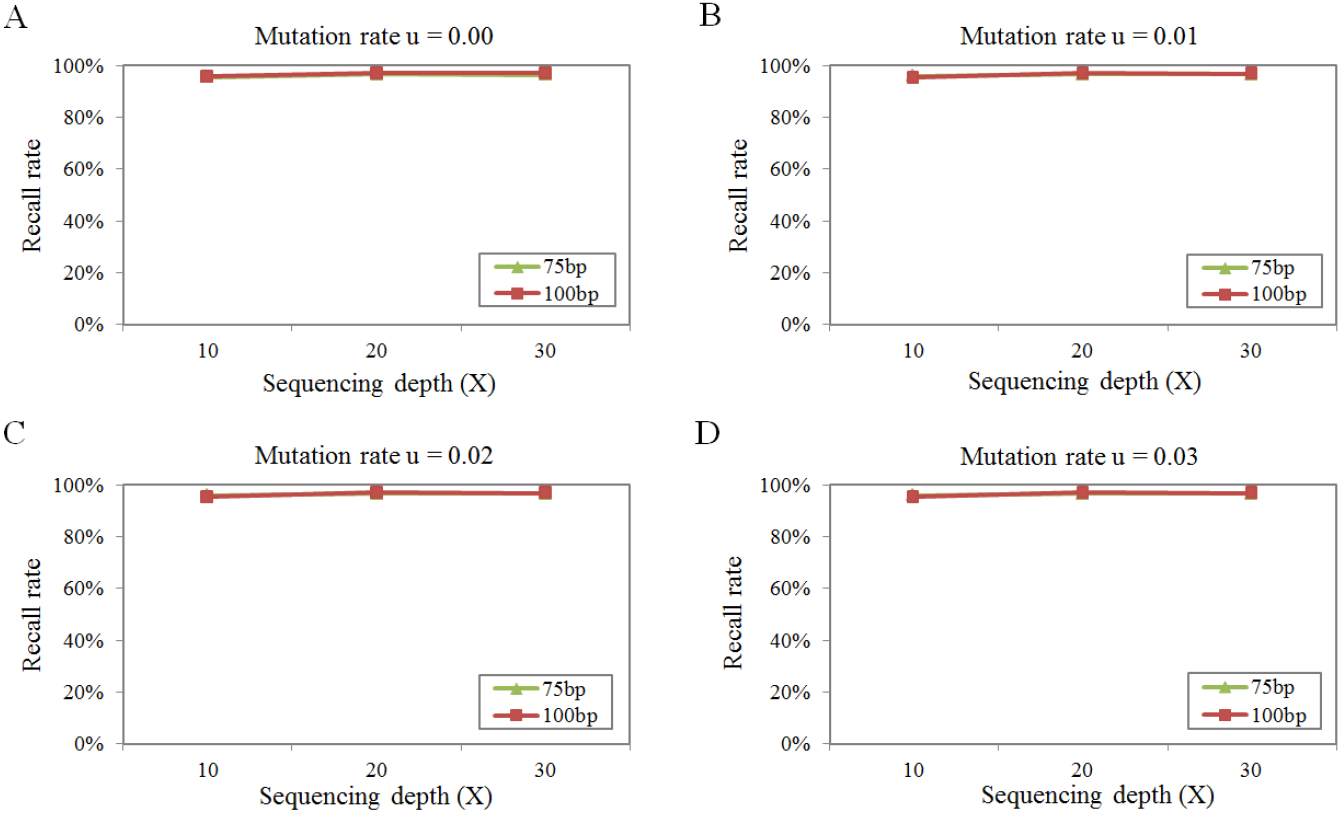
Recall rates of the discovery of retroposed parental genes with optimized parameter settings. The optimized discovery parameter setting for each simulated genomic sequencing dataset is provided in the bottom panel of *SI Appendix* Fig. S3. The recall rate for each simulated genomic sequencing dataset was calculated as the fraction of identified ‘true’ retroposed parental genes from 1,000 random simulations.

**Fig. S6.**
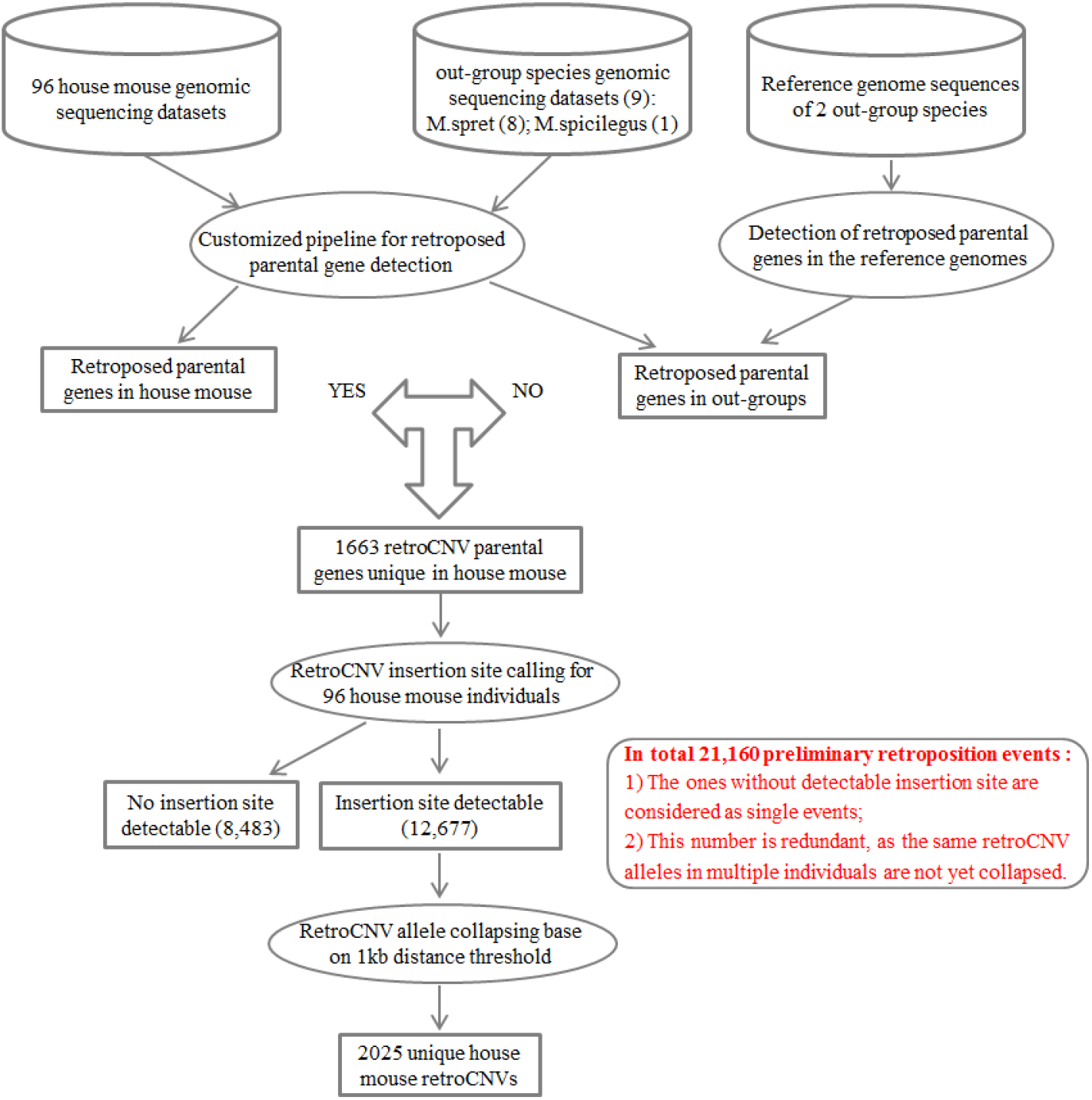
A flow chart to show the detection of house mouse unique retroCNVs. With the above customized pipeline for retroposed parental gene detection, we detected the retroposed parental genes for both 96 house mouse individuals and 9 out-group species mice individuals based on the genomic sequencing datasets. Additionally, we also obtained another set of retroposed parental genes for 2 out-group species, based on their reference genome sequences (See Materials and Methods). The 1663 retroposed parental genes that are present in the house mouse individuals but absent in the out-group species were taken as the retroCNV parental genes in house mouse. Insertion site calling on these retroCNV parental genes returned 21,160 preliminary retroposition events in 96 house mouse individuals. For those ones with insertion site detectable (12,667), we further collapsed the same retroCNV alleles in multiple individuals with 1kb distance clustering threshold. Finally, we obtained 2025 unique house mouse retroCNVs for further analysis.

**Fig. S7.**
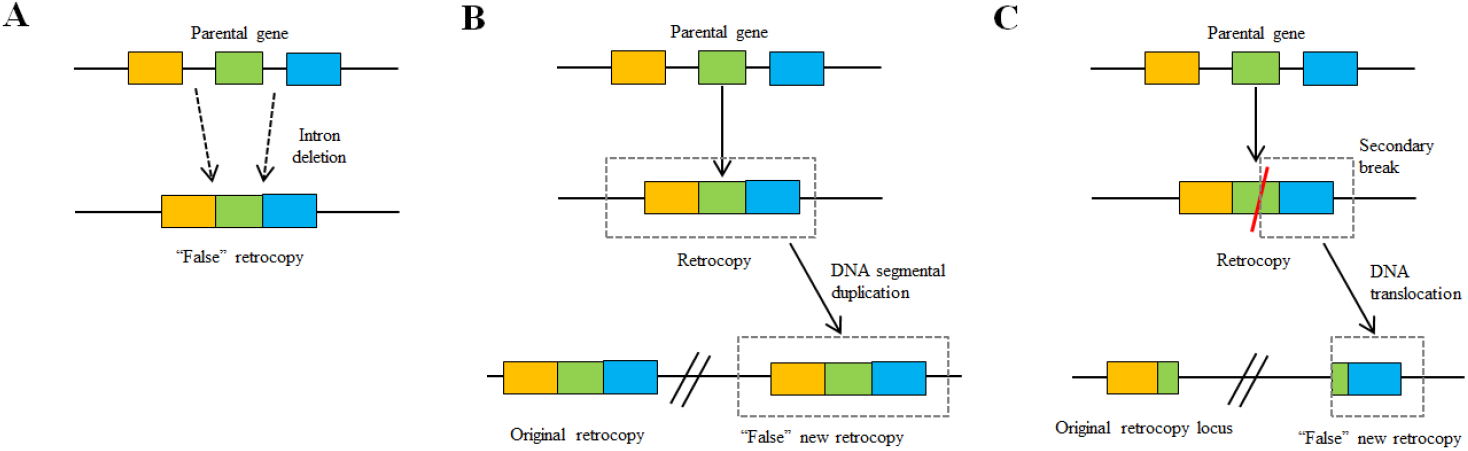
Possible scenarios to cause false discovery of retrocopies. A) Intron deletion event to generate retrocopy alike intron-free gene structure; B) DNA segmentation of existing retrocopy to generate an additional “false” new retrocopy; C) Secondary break of existing retrocopy to generate “false” new retrocopy.

**Fig. S8.**
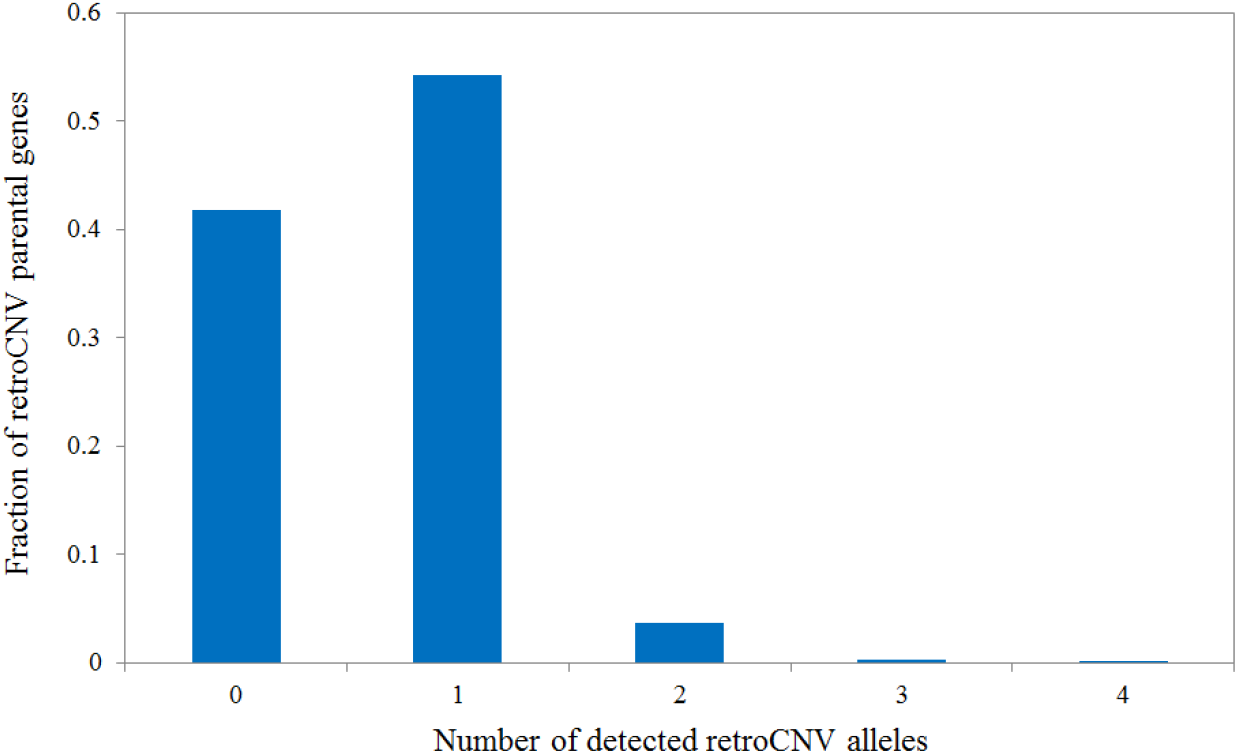
Distribution of the number of detected retroCNV alleles for retroCNV parental genes. The calculation on the number of retroCNV alleles for each retroCNV parental gene was based on the data provided in Dataset S2. It was based on the number of retrocopies (present in the mm10 reference genome) or insertion sites (full-length retrocopies absent in the mm10 reference genome) found in the genome of a given animal. 0 means that the insertion site for a given retroCNV parental gene could not be uniquely localized likely due to having landed in a repetitive genome region.

**Fig. S9.**
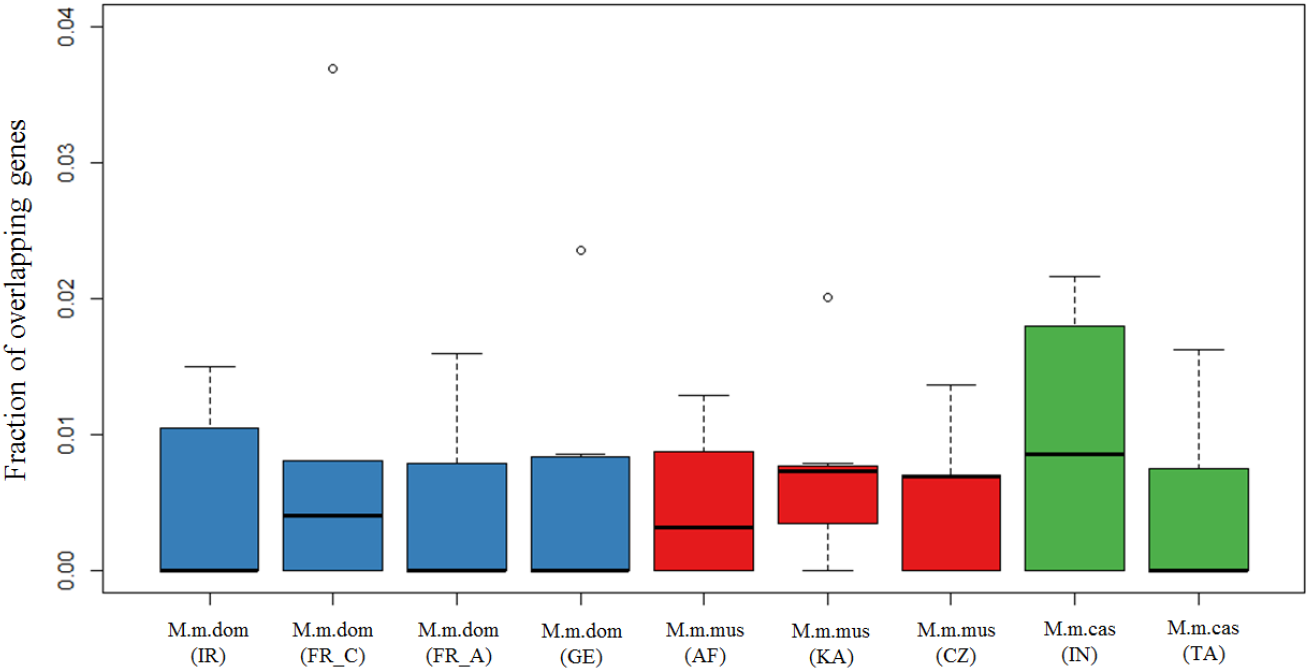
Fraction of overlap between retroCNV parental genes and genic CNVs across house mouse populations. The fraction of overlap for each individual was defined as the number of overlapping genes divided by the average number of detected retroCNV parental genes and genic CNVs. Abbreviations for geographic regions: IR, Iran; FR_C, France (Central Massif); FR_A, France (Auvergne-Rhône-Alpes); GE, Germany; AF, Afghanistan; KA, Kazakhstan; CZ, Czech Republic; IN, India; TA, Taiwan.

**Fig. S10.**
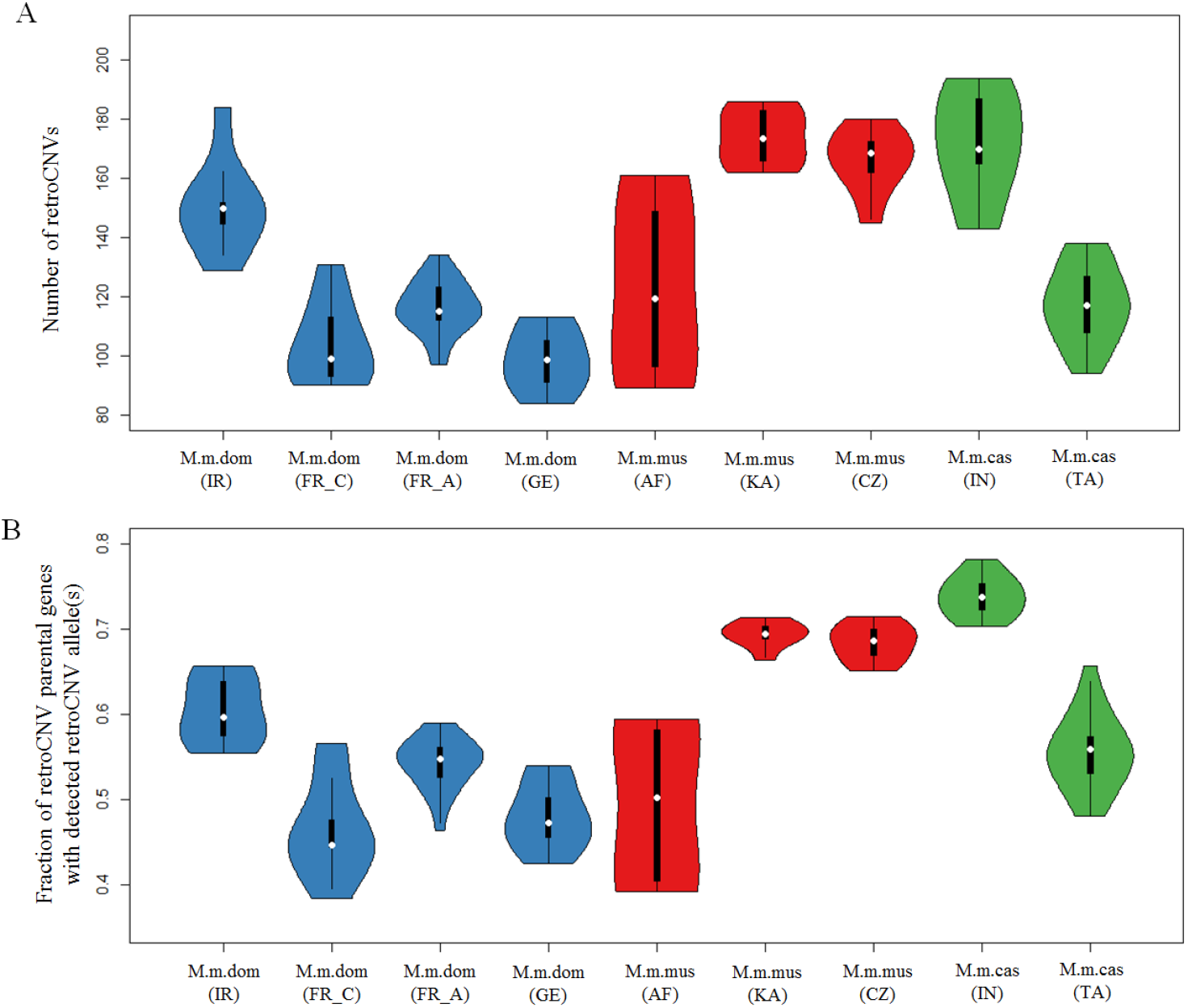
Distribution of gene retroposition events across house mouse natural populations. (A) Distribution of number of retroCNVs among different house mouse natural populations. In case of multiple insertion sites (or retrocopies present in the mm10 reference genome) detected for one retroCNV parental gene, each insertion site was taken as one independent gene retroposition event (*i.e*., retroCNV). Note that Fig. 2B in the main text shows the corresponding numbers for retroCNV parental genes. The overall pattern is similar, but with a relatively lower discovery rate in the Afghanistan population, which likely attributes to the shorter read length and insert size of the sequencing data for individuals from this population (*SI Appendix*, Table S1). (B) shows the fraction of retroCNV parental genes with detectable retroCNV allele(s) of individuals across nine house mouse natural populations. The calculation on the fraction of detectable retroCNV allele(s) was based on the data provided in Dataset S2. The abbreviations for geographic regions follow Fig. 1 / *SI Appendix* Fig. S9.

**Fig. S11:**
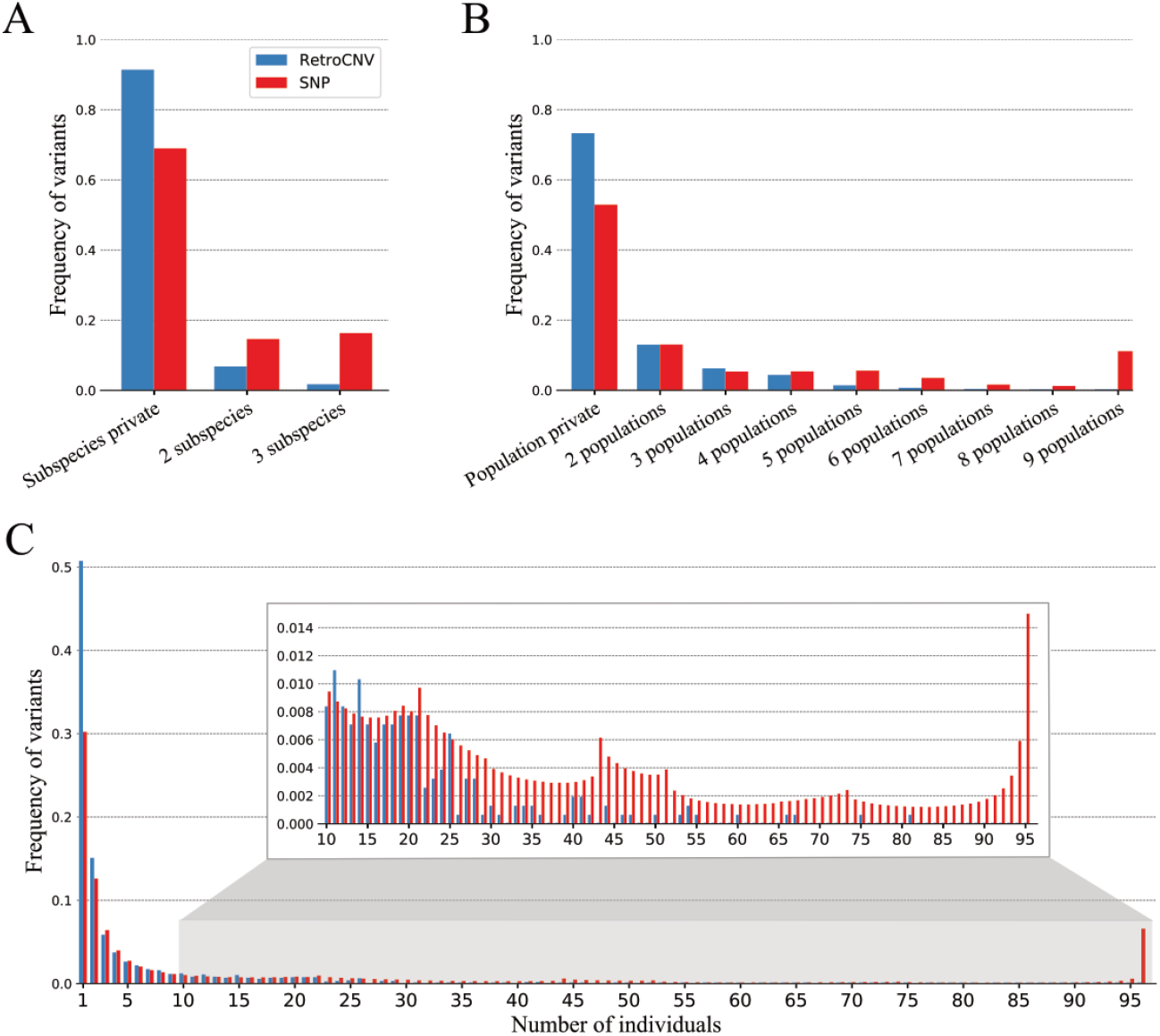
Distribution of the allele frequency of retroCNVs and SNPs. Complementary to Fig. 3 in the main text, this figure represents an additional analysis with only retroCNVs that show both positive evidence of retroCNV presence (*i.e*., detectable retroCNV allele) and positive evidence of retroCNV absence (*i.e*., alignments to span the insertion site of retroCNV) in all 96 tested house mouse individuals. Enlarged in the inset box is to show the frequencies of retroCNVs/SNPs present in larger number of individuals.

**Fig. S12:**
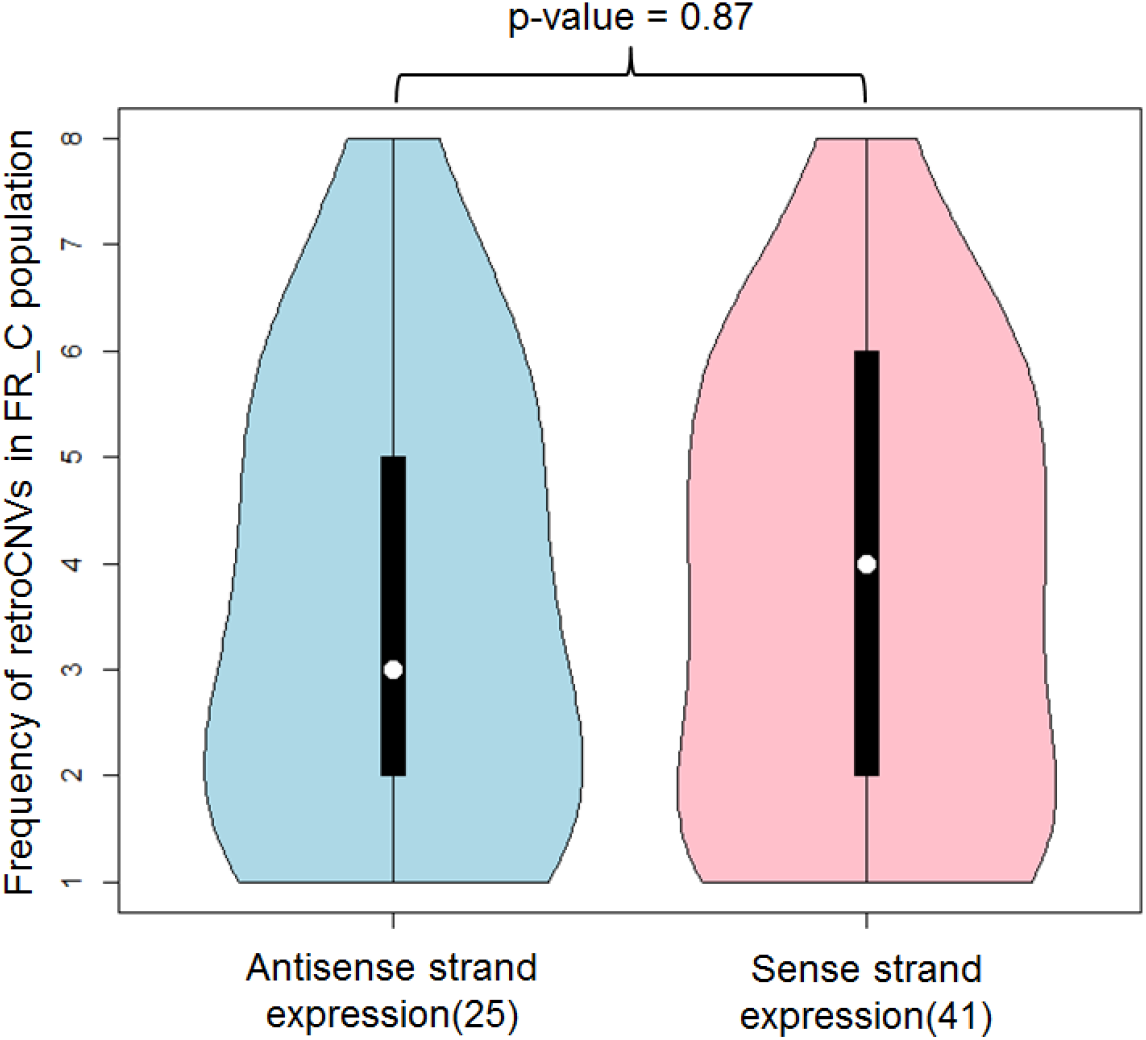
Comparison of allele frequency of retroCNVs transcribed on the antisense strand and sense strand. As the strand-specific RNA-seq dataset was generated from mice individuals of the FR_C population, the allele frequencies of retroCNVs were computed based on only the same population individuals. The expression of a retroCNV is defined as FPKM >0 where at least one read could be uniquely mapped to retroCNV (23). The number of retroCNVs with expression (in at least 1 tissue) for each category is listed in the parentheses after each. The statistical significance was calculated by using Wilcoxon rank sum test.

**Table S1.**
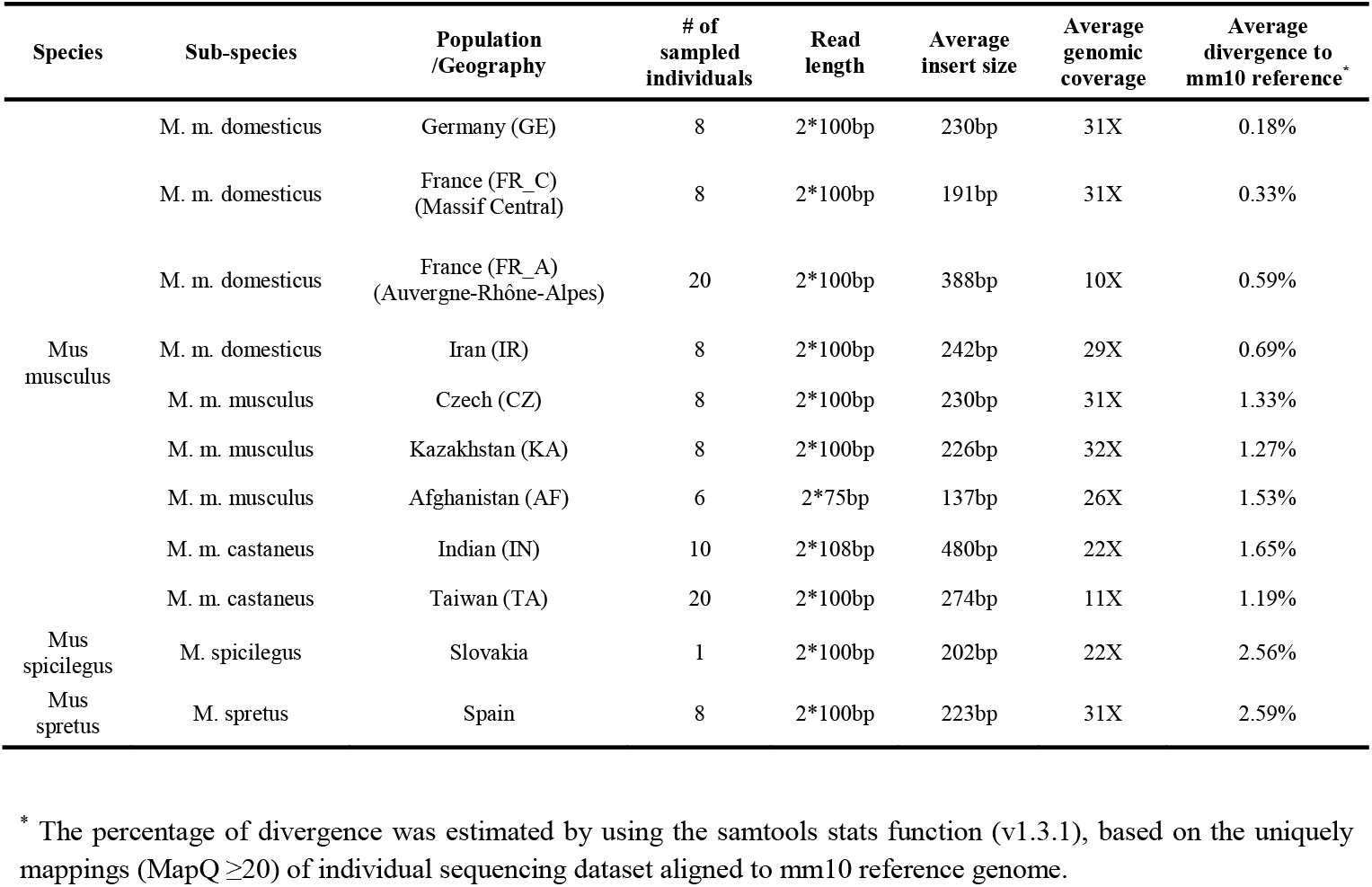
Summary statistics of genomic short read sequencing datasets

**Table S2.** Summary of RetroCNV expression in the non-strand-specific RNA-Seq dataset

|  |  | Testis | Brain | Gut | Muscle | Kidney | Spleen | Liver | Heart | Lung | Thyroid |
| --- | --- | --- | --- | --- | --- | --- | --- | --- | --- | --- | --- |
| M.m.dom_GE | # of expressed retroCNVs (FPKM>0) | 40 | 29 | 20 | 24 | 31 | 33 | 26 | 25 | 28 | 28 |
|  | <sup>1</sup>Average expression level in FPKM | 3.1<br>(<sup>2</sup>SEM:1.1) | 2.9<br>(SEM:1.0) | 3.1<br>(SEM:1.2) | 3.5<br>(SEM:1.8) | 3.1<br>(SEM:1.1) | 1.9<br>(SEM:0.7) | 1.9<br>(SEM:0.6) | 4.7<br>(SEM:2.0) | 2.1<br>(SEM:0.7) | 1.7<br>(SEM:0.6) |
| M.m.dom_FR_C | # of expressed retroCNVs (FPKM>0) | 32 | 21 | 20 | 16 | 28 | 25 | 18 | 18 | 17 | 20 |
|  | <sup>1</sup>Average expression level in FPKM | 3.7<br>(SEM:1.6) | 2.4<br>(SEM:0.7) | 2.2<br>(SEM:0.7) | 3.1<br>(SEM:1.5) | 2.2<br>(SEM:0.7) | 1.8<br>(SEM:0.6) | 2.0<br>(SEM:0.5) | 3.6<br>(SEM:1.6) | 1.8<br>(SEM:0.7) | 1.6<br>(SEM:0.5) |
| M.m.dom_IR | # of expressed retroCNVs (FPKM>0) | 36 | 20 | 22 | 15 | 23 | 18 | 21 | 22 | 24 | 24 |
|  | <sup>1</sup>Average expression level in FPKM | 2.4<br>(SEM:0.7) | 3.0<br>(SEM:1.0) | 1.9<br>(SEM:0.7) | 3.6<br>(SEM:1.7) | 4.2<br>(SEM:1.7) | 1.0<br>(SEM:0.3) | 1.8<br>(SEM:0.6) | 4.7<br>(SEM:2.3) | 1.5<br>(SEM:0.4) | 0.8<br>(SEM:0.2) |

<sup>1</sup> Only the retroCNVs with non-zero expression were included;
SEM: standard error of mean.

### Legends for Datasets S1 to S6

**Dataset S1** List of genomic and transcriptomic sequencing datasets used in this study

**Dataset S2** List of gene retroposition events detected in all house mouse individuals

**Dataset S3** List of all house mouse specific retroCNVs

**Dataset S4** RetroCNV expression based on the non-strand specific RNA-Seq dataset

**Dataset S5** The impact on parental gene expression for singleton retroCNVs

**Dataset S6** RetroCNV expression on the sense/antisense strand based on strand-specific RNA-Seq dataset

